# Brain Network Organization During Mindful Acceptance of Emotions

**DOI:** 10.1101/2020.03.31.018697

**Authors:** Matthew Luke Dixon, Manesh Girn, Kalina Christoff

## Abstract

Individuals use various strategies to cope with challenging emotions such as anxiety. Mindful acceptance involves broadening attentional scope and fully experiencing present moment sensory feelings (whether pleasant or unpleasant) without judgment or elaboration. In contrast, narrative-evaluation involves focusing on a narrow band of sensory experience and generating an elaborate narrative about the meaning and desirability of one’s emotional feelings. The current study examined brain network organization during these strategies using graph theoretic analyses. We used a naturalistic task paradigm in which participants reflected on an anxiety-provoking issue from their personal lives and adopted each strategy in different blocks. Compared to narrative-evaluation, mindful acceptance was associated with: (i) increased global network connectivity; (ii) greater integration of interoceptive regions (mid and posterior insula) into large-scale networks; (iii) reorganization of motivational circuits including a shift in the striatum’s network assignment from the default network to the salience network; and (iv) a shift from default network to frontoparietal control network (FPCN) regions as central hubs that coordinate information flow. Functional connectivity patterns within the left FPCN were associated with acceptance reports. Thus, broadening attentional scope during mindful acceptance is supported by a more globally interconnected neural landscape, as well as greater information flow through FPCN regions that underlie metacognitive awareness and cognitive control.

---

> “Flow with whatever may happen, and let your mind be free.”
>
> — — Zhuangzi, Nan-Hua-Ch’en-Ching

## Introduction

There is considerable interest in identifying effective strategies for handling challenging emotions such as anxiety. The importance of this endeavour is underscored by the personal and societal costs of affective disorders. Over the past two decades there has been rigorous scientific investigation into strategies for altering the nature, duration, and intensity of emotions (Gross, 1998, 2015; Ochsner, Silvers, & Buhle, 2012; Webb, Miles, & Sheeran, 2012). While considerable work has focused on strategies such as distraction and cognitive reappraisal, there is growing interest in a distinct form of emotion regulation known as mindful acceptance of emotions. Acceptance is one of several components of mindfulness practice (Baer, Smith, Hopkins, Krietemeyer, & Toney, 2006; Bishop et al., 2004; Britton et al., 2018; Holzel et al., 2011; Lindsay & Creswell, 2017) and has its roots in contemplative traditions including Buddhism (Harvey, 1990). A guiding principle underlying mindful acceptance is the notion that a source of unhappiness is resistance to present moment experience. While most individuals feel a desire to avoid so called “negative” emotions, this desire itself can paradoxically cause suffering given that these emotions will naturally arise from time to time in response to life challenges.

The aim of mindful acceptance is to fully experience emotional feelings ‒ whether pleasant or unpleasant ‒ without trying to control, interpret, change, or avoid them. It involves staying fully conscious with present moment sensory experience by adopting an open and non-judgmental stance towards inner thoughts, emotions, and sensations, and disengaging from elaborative thinking, judgement, or defense mechanisms (Farb, Anderson, & Segal, 2012a; Hayes, Luoma, Bond, Masuda, & Lillis, 2006; Kabat-Zinn, 1994). Thus, during the experience of challenging emotions, mindful acceptance involves a broadening of attention scope (Farb, et al., 2012a) and fully registering the concrete aspects of uncomfortable feelings (e.g., raw sensations such as muscle tension, the urge to withdraw, shallow breathing, and also more complex feeling states such as heartache). While it seems counterintuitive that more deeply experiencing uncomfortable feelings can enhance well-being, acceptance has been shown to reduce cognitive and physiological reactivity, encourage free-flowing mental activity and insight, increase behavioral expressivity, and foster resilience through exposure to feared emotional sensations (Arch & Craske, 2006; Campbell-Sills, Barlow, Brown, & Hofmann, 2006; Chambers, Gullone, & Allen, 2009; Dan-Glauser & Gross, 2015; Farb et al., 2010; Kabat-Zinn, 2009; Lindsay, Young, Smyth, Brown, & Creswell, 2018; Segal, Williams, & Teasdale, 2013). Notably, acceptance does not entail a complete suppression of preferences and meaning. Rather, it is about limiting habitual reactions, interpretations, and excessive elaboration of sensory processing, which may in fact create space for novel insights and positive appraisals to arise (Farb, et al., 2012a; Garland, Farb, Goldin, & Fredrickson, 2015). The idea of working with, rather than fighting against, present moment experience has been incorporated into a variety of approaches aimed at boosting psychological well-being and treating affective disorders (Britton, Shahar, Szepsenwol, & Jacobs, 2012; Farb, et al., 2012a; Goldin et al., 2016; Hayes, et al., 2006; Hofmann, Sawyer, Witt, & Oh, 2010; Kabat-Zinn, 1994; Kuyken et al., 2015; Linehan, 1993; Segal, Williams, & Teasdale, 2002).

At the opposite end of the spectrum is a narrative-evaluative mindset, which often arises spontaneously during the experience of uncomfortable emotions. This mindset involves narrowing attention around a limited band of sensory input and engaging in elaborative self-referential thinking in an attempt to reconcile one’s current experience with one’s an autobiographical narrative – a temporally extended story about ‘me’ (Damasio, 1998; Gallagher, 2000). This narrative-evaluative mode of processing often involves resisting and judging uncomfortable feelings as bad and the desire to escape those feelings. Although people generally assume that thinking about their emotional experience is useful and will help them solve the problems that gave rise to those feelings, it may in some cases result in repetitive loops of maladaptive thinking that exacerbate negative affect and lead the individual further and further away from present moment sensory experience (Nolen-Hoeksema, Wisco, & Lyubomirsky, 2008). Just as individuals can direct attention to modulate sensory processing to emphasize one object or spatial location over another (Brefczynski & DeYoe, 1999; Buschman & Kastner, 2015; Desimone & Duncan, 1995; Kanwisher & Wojciulik, 2000), so too can they adjust the scope of attention to emphasize sensory-based experiencing or narrative-evaluative processing during an emotional response.

Neuroscientific studies have begun to delineate the neural circuitry that may underlie these distinct ways of attending to emotional experience. The insula has received considerable attention in the context of interoception and mindfulness. In particular, the mid and posterior insula are thought to be primary cortical interoceptive regions that generate a map of bodily signals arriving via the lamina I spinothalamic and vagus nerve homeostatic pathways (Barrett & Simmons, 2015; Craig, 2002; Critchley & Harrison, 2013; Farb, Segal, & Anderson, 2012b). These signals are combined with broader contextual information in the anterior insula, giving rise to complex subjective feelings (Craig, 2002). Prior work has shown that insula activation increases when participants adopt a sensory-based experiential mindset compared to a narrative-based mindset during self-reflection (Farb et al., 2007). Furthermore, insula activation correlates with interoceptive accuracy when participants monitor their heartbeat (Critchley, Wiens, Rotshtein, Ohman, & Dolan, 2004). Notably, meditation training is associated with greater interoceptive accuracy (Fox et al., 2012) and is associated with changes in insula structure and function (Farb, Segal, & Anderson, 2013; Fox et al., 2014; Young et al., 2017). The mid and posterior insula have a close relationship with sensorimotor cortices, which may also play a role in representing the concrete aspects of sensory experience during emotional processing (Khalsa, Rudrauf, Feinstein, & Tranel, 2009; Satpute et al., 2015).

On the other hand, the default network sits at the top of the cortical processing hierarchy (Margulies et al., 2016) and plays a central role in self-referential processing and evaluation. Activation in default network regions is associated with narrative-based self-reflection (Farb, et al., 2007), autobiographical memory retrieval and future planning (Andrews-Hanna, Saxe, & Yarkoni, 2014a; Spreng, Stevens, Chamberlain, Gilmore, & Schacter, 2010), reflection on personal values (D’Argembeau et al., 2012), self-referential evaluations (Dixon, Thiruchselvam, Todd, & Christoff, 2017c), inferring mental states (Andrews-Hanna, et al., 2014a; Spreng, Mar, & Kim, 2009; Van Overwalle, 2009), spontaneous thought (Christoff, Gordon, Smallwood, Smith, & Schooler, 2009a; Fox, Spreng, Ellamil, Andrews-Hanna, & Christoff, 2015), and thinking about why actions are performed (Spunt, Falk, & Lieberman, 2010). Thus, default network regions may place emotional signals in the context of a temporally-extended personal narrative that includes cognitive elaborations, personal meaning, and preferences (Andrews-Hanna, Smallwood, & Spreng, 2014b; Dixon, et al., 2017c; Farb, et al., 2007; Fox et al., 2018). Moreover, the core subsystem of the default network is thought to constrain the flow of thoughts based on affective meaning (Christoff, Irving, Fox, Spreng, & Andrews-Hanna, 2016). In line with this, a growing literature suggests that elevated activation and altered connectivity patterns of default network regions may contribute to ruminative tendencies and clinical disorders (Berman et al., 2011; Dixon et al., 2020; Hamilton, Farmer, Fogelman, & Gotlib, 2015; Kaiser, Andrews-Hanna, Wager, & Pizzagalli, 2015; Kross, Davidson, Weber, & Ochsner, 2009; Kucyi et al., 2014; Ray et al., 2005). Finally, studies have demonstrated that meditation training is associated with the capacity to deactivate this network (Brewer et al., 2011; Farb, et al., 2007; Garrison et al., 2013b). Together, these findings suggest a distinction between insular and sensorimotor cortices that support sensory-based experiencing, and default network regions that support self-referential meaning and evaluation.

How does the brain switch between acceptance of sensory-based experiencing and narrative-evaluative processing? Several lines of evidence suggest that the frontoparietal control network (FPCN) may play a critical role in this process. The FPCN contributes to cognitive control and controlling the focus of attention through its role in working memory (Bunge, Ochsner, Desmond, Glover, & Gabrieli, 2001; D’Esposito & Postle, 2015; Owen, McMillan, Laird, & Bullmore, 2005), representing task sets and rules (Bunge, Kahn, Wallis, Miller, & Wagner, 2003; Dixon & Christoff, 2012; Dixon, Girn, & Christoff, 2017b; Duncan, 2013; Koechlin, Ody, & Kouneiher, 2003; Sakai, 2008), higher-order (abstract) thought and reasoning (Christoff, Keramatian, Gordon, Smith, & Madler, 2009b; Christoff et al., 2001), and metacognitive awareness (Fleming, Weil, Nagy, Dolan, & Rees, 2010; McCaig, Dixon, Keramatian, Liu, & Christoff, 2011). The FPCN is thought to modulate activity in other brain networks based on current goals and task demands (Cole, Repovs, & Anticevic, 2014b; Fornito, Harrison, Zalesky, & Simons, 2012; Spreng, et al., 2010) through its widespread connectivity (Cole et al., 2013; Dixon et al., 2018; Spreng, Sepulcre, Turner, Stevens, & Schacter, 2013). Moreover, the FPCN’s functional connectivity profile reconfigures across a variety of task states to support the flexible implementation of specific task demands (Cole, et al., 2013). By exerting top-down control over other networks, the FPCN may regulate attention and the extent to which sensory information versus higher-order thought is emphasized (Dixon, et al., 2018; Dixon, Fox, & Christoff, 2014). In the context of mindful acceptance versus narrative-evaluative processing, this may occur through influencing the nature of processing in the insula and default network.

Despite the progress that has been made in understanding the neural basis of mindful acceptance and how it differs from other emotion regulation strategies (e.g., reappraisal) (Farb, et al., 2010; Farb, et al., 2013; Farb, et al., 2007; Goldin, Jazaieri, & Gross, 2014; Goldin, Ziv, Jazaieri, Hahn, & Gross, 2013; Goldin, Moodie, & Gross, 2019; Holzel, et al., 2011; Lutz, Slagter, Dunne, & Davidson, 2008; Smoski et al., 2015; Tang, Holzel, & Posner, 2015; Vago & Zeidan, 2016), we currently know little about brain network organization during acceptance and how it differs from narrative-evaluative processing. A network neuroscience approach conceptualizes the brain as a set of nodes (regions) and connections, and provides a way to summarize the multivariate relationships that underlie complex cognitive processes (Cole, Ito, Bassett, & Schultz, 2016; Fornito, Zalesky, & Bullmore, 2016; Medaglia, Lynall, & Bassett, 2015; Petersen & Sporns, 2015; Rubinov & Sporns, 2010). This approach may offer valuable information about the neural basis of introspective processes, because they likely rely on interactions across distributed cortical and subcortical regions. While there is some evidence of a stable network backbone (Cole, Bassett, Power, Braver, & Petersen, 2014a; Krienen, Yeo, & Buckner, 2014), there is also growing evidence that network properties dynamically change across time and task context (Cohen & D’Esposito, 2016; Gonzalez-Castillo & Bandettini, 2017; Hutchison et al., 2013; Medaglia, et al., 2015; Metzak et al., 2011; Shine et al., 2016). Studies of task-related changes in functional connectivity have mainly used traditional cognitive tasks that involve responding to externally presented stimuli that lack personal relevance (e.g., comparing network configuration during an *N*-back working memory versus rest). Thus, little is currently known about the capacity for network reorganization during naturalistic tasks, including introspective states that involve self-relevant and emotionally-charged content. Studies have begun to examine emotional processes from a network perspective (Barrett & Satpute, 2013; Brandl et al., 2018; Hermans et al., 2011; McMenamin, Langeslag, Sirbu, Padmala, & Pessoa, 2014; Pessoa, 2017), but have yet to address how network properties change when a mindful acceptance or narrative-evaluative mindset is applied to emotional experience. Additionally, while numerous studies have examined network properties in relation to meditation training (Brewer, et al., 2011; Creswell et al., 2016; Farb, et al., 2013; Farb, et al., 2007; Hasenkamp & Barsalou, 2012; Jang et al., 2011; Jao et al., 2016; Josipovic, Dinstein, Weber, & Heeger, 2011; Pagnoni, 2012; Tang, Tang, Tang, & Lewis-Peacock, 2017; Taren et al., 2015; Xue, Tang, & Posner, 2011), few studies have specifically targeted emotional processes or compared network organization during different types of introspective states within the same individuals.

In the current study, we examined brain network organization during mindful acceptance and narrative-evaluation in participants that were not pre-selected based on meditation experience. We used a naturalistic task paradigm in which healthy participants were asked to reflect upon an anxiety-provoking interpersonal situation from their life that was currently unresolved. In separate fMRI scans, participants reflected on this issue while adopting a narrative-evaluation mindset or an acceptance mindset based on detailed instructions provided before scanning (see **Methods**). We used graph theory to examine global and local network properties (**Figure 1**). First, we used a community detection algorithm to create a data-driven network parcellation. This allowed us to examine whether nodes group into networks in different ways across conditions. Second, we examined the capacity for specialized modular processing using measures of clustering and system segregation. This allowed us to look for condition-dependent changes in more fine-grained or local network properties. Third, we examined potential changes across conditions in global connectivity and the ease of information transfer across the whole network using a measure known as global efficiency. Fourth, we quantified the extent to which each node acted as a ‘hub’ that facilitates information flow and integration and whether the nodes identified as hubs differed across conditions. Finally, we examined the relationship between multivariate FC patterns and individual variation in subjective experience – that is, subjective reports about the quality of mindful acceptance during the task.

**Figure 1.**
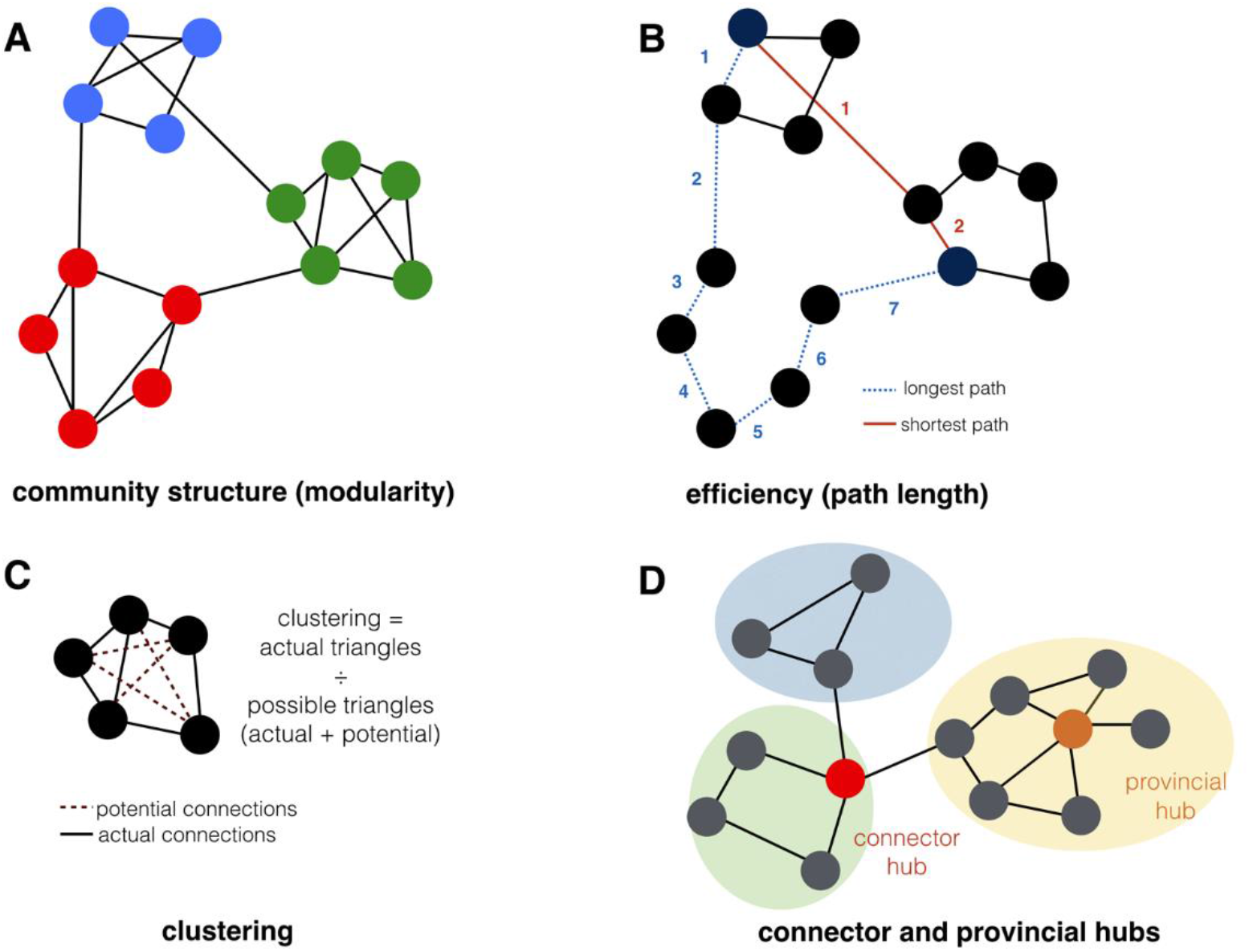
Basic graph theory concepts. **(A)** Example of community structure in a modular network with three distinct communities or networks containing stronger within-network than between-network connections. **(B)** Global efficiency relates to path length between nodes. The blue dotted path requires seven edges to connect targets α and β, compared to the solid red path which only requires two edges. Information transfer through the red path would be more efficient, and therefore, networks containing more of such short pathways have higher global efficiency. **(C)** Example of clustering within a network, which provides an index of the strength of communication within a local neighborhood of nodes. Dashed lines represent possible connections. Solid lines represent actual connections. When three nodes are fully connected, a triangle forms, representing a densely interconnected neighborhood. Clustering is computed based on the number of actual triangles divided by the number of possible triangles in a network. Here only one triangle is actually formed out of all possible triangles, demonstrating low clustering within this set of nodes. **(D)** Some nodes act as central hubs that support information integration and distribution and exert strong influence on overall network dynamics by virtue of having widely distributed connections. Connector hubs have distributed connections across multiple communities, whereas provincial hubs have distributed connections within their home community. Figure adapted from (Girn, Mills, & Christoff, 2019).

## METHODS

### Participants

Participants were 24 healthy adults (Mean age = 30.33, SD = 4.80; 10 female; 22 right handed) living in the greater Vancouver area, with no history of head trauma or psychological conditions. This study was approved by the University of British Columbia clinical research ethics board, and all participants provided written informed consent, and received payment ($20/hour) for their participation. Due to a technical error, fMRI data for the acceptance condition was not collected for one participant, and fMRI data for 3 other participants was excluded due to excessive movement (described below), resulting in a final fMRI data sample of 20 participants. At the end of scanning, one participant reported experiencing physical discomfort throughout the scan. Similar results were obtained with or without inclusion of this participant’s data, so they were included in the final analysis. Participants were not selected based on prior meditation experience, but had a wide range of total (lifetime) hours of meditation practice (Median = 40; range: 0 - 2366), consistent with the ease of access and popularity of yoga and meditation in Vancouver. The data used in this study is part of a larger data set (Dixon et al., 2017a; Dixon, et al., 2018).

### Experimental task conditions

Participants performed an autobiographical emotion provocation task in which they were asked to think about a specific person in their life (e.g., a friend, roommate, sibling, or partner) and to reflect on an aspect of the person’s personality or an interaction with them that was mildly upsetting and anxiety-provoking, and was a currently unresolved issue that they wished was different. Participants were instructed to hold this scenario in mind whilst employing each mindset in separate six-minute scans. Prior to scanning, participants underwent a training procedure to ensure that they understood how to implement the two mindsets. Instructions were delivered by an author (MLD) with over 16 years of meditation experience (at the time of testing), including 12 residential retreats. Participants were not instructed to alter their emotional experience directly when employing each mindset. The instructions were repeated just prior to each scan. Participants were asked to select the person/situation that they would think about before the scanning session began. The condition order was fixed, with the narrative-evaluation condition always preceding the acceptance condition. This was done because we hypothesized that the acceptance condition would be more difficult to implement and that it may have a lasting impact and bleed-over into the narrative-evaluation condition if it were to be performed first.

#### Narrative-evaluative mindset

We asked participants to adopt a narrative-evaluative mindset for the duration of the 6-minute run with the following instructions: “As you think about this person and situation, reflect on why this situation is upsetting to you. Analyze whether you like or dislike the feelings associated with this situation. Think about why the person is the way they are; who caused the situation and why; what has happened in the past to lead up to this point in your relationship; and what might happen going forwards into the future ― how things might get worse or better. Think about how you would feel if things with this person were different. As you think about this topic, allow yourself to become fully caught up in your thoughts and emotions. Try to think about what the person and situation means to you, and what aspects are good or bad.” This type of mindset thus emphasized: (i) a temporally extended episode; (ii) judgment of feelings as good or bad; and (iii) analysis of the meaning and cause/consequences of feelings. Importantly, we did not ask participants to change their mood or feelings in any way. Participants were instructed to keep their eyes closed, but were told that they could open their eyes if they needed a reminder of the instructions (short sentences capturing the essence of the instructions were presented on screen). Participants were asked to provide a brief description of their understanding of the instructions to ensure that they knew how to adopt the mindset.

#### Acceptance mindset

In this condition, participants were asked to reflect on the same upsetting issue as in the case of the narrative-evaluative condition, but this time, received the following instructions: “In this condition, hold that same person/situation in mind, but do your best to avoid any type of mental analysis of the situation. Instead, pay attention to the feelings of your body in the present moment. For example, as you think of this person, notice whether your heart starts to beat faster, or whether your body becomes tense or relaxed, and notice how you are breathing. Try your best to stay with the feelings of your body as they come and go. When thoughts or feelings arise, try to observe them and then let them go. Think of your feelings as waves that emerge and then disappear back into the ocean. Try not to analyze your thoughts and feelings as good or bad, and try not to think about the cause of the situation or what might happen in the future. Just keep returning to the sensations of your body in the present moment, and allow yourself to accept and experience them fully, whatever they are. Whenever you notice that you have become caught up in a train of thought or start to analyze your experience, that’s okay; just calmly re-focus on how your body is feeling in the present moment. There is no specific goal or purpose other than noticing what you are thinking and feeling from one moment to the next.” This mindset thus emphasized: (i) present moment focus on the concrete sensations of the body; (ii) non-judgment of feelings; and (iii) letting go of analysis related to the meaning and cause/consequences of feelings. Importantly, we did not use the term mindfulness when describing this condition, given its positive connotations, nor did we instruct participants that this type of mindset may improve mood. They were not asked to change their mood or feelings in any way. Participants were instructed to keep their eyes closed, but were told that they could open their eyes if they needed a reminder of the instructions. Participants were asked to provide a brief description of their understanding of the instructions to ensure that they knew how to adopt the mindset.

#### Self-reports

Following the scanning session, participants filled out a questionnaire pertaining to their experience during the conditions. Participants rated the following questions on a scale from 1 = not at all, to 7 = a lot / very much: (1) To what extent were you aware of your body and experiencing bodily sensations? (2) To what extent do you feel like you were caught up in your thoughts and feelings? (3) To what extent were you focused on analyzing the causes of the situation? (4) To what extent did you think about what might happen in the future in relation to this situation? (5) To what extent did you judge your thoughts/feelings as good or bad? (6) How difficult was it to follow the instructions and evaluate/accept your thoughts and feelings? Participants also rated their mood during the condition on a 7-point scale from 1 = very unpleasant to 7 = very pleasant. Participants also reported the number of hours they had spent engaged in meditation in the past.

#### Resting state

Prior to the narrative-evaluative and acceptance conditions, there was a resting state scan during which participants lay in the scanner with their eyes closed and were instructed to relax and stay awake, and to allow their thoughts to flow naturally. This was used as a comparison condition for the network-based parcellation analysis.

### MRI data acquisition

fMRI data were collected using a 3.0-Tesla Philips Intera MRI scanner (Best, Netherlands) with an 8-channel phased array head coil with parallel imaging capability (SENSE). Head movement was restricted using foam padding around the head. T2*-weighted functional images were acquired parallel to the anterior commissure/posterior commissure (AC/PC) line using a single shot gradient echo-planar sequence (repetition time, TR = 2 s; TE = 30 ms; flip angle, FA = 90°; field of view, FOV = 240 mm; matrix size = 80 × 80; SENSE factor = 1.0). Thirty-six interleaved axial slices covering the whole brain were acquired (3-mm thick with 1-mm skip). Each session was six minutes in length, during which 180 functional volumes were acquired. Data collected during the first 4 TRs were discarded to allow for T1 equilibration effects. Before functional imaging, a high resolution T1-weighted structural image was acquired (170 axial slices; TR = 7.7 ms; TE = 3.6 ms; FOV = 256 mm; matrix size = 256 × 256; voxel size = 1 × 1 × 1 mm; FA = 8°). Total scan time was ~ 60 minutes. Head motion was minimized using a pillow, and scanner noise was minimized with earplugs.

### Preprocessing

Image preprocessing and analysis were conducted with Statistical Parametric Mapping (SPM8, University College London, London, UK; http://www.fil.ion.ucl.ac.uk/spm/software/spm8). The time-series data were slice-time corrected (to the middle slice), realigned to the first volume to correct for between-scan motion (using a 6-parameter rigid body transformation), and coregistered with the T1-weighted structural image. The T1 image was bias-corrected and segmented using template (ICBM) tissue probability maps for gray/white matter and CSF. Parameters obtained from this step were subsequently applied to the functional (re-sampled to 3 mm^3^ voxels) and structural (re-sampled to 1 mm^3^ voxels) data during normalization to MNI space using 4^th^ order b-spline interpolation. The data were spatially-smoothed using an 8-mm^3^ full-width at half-maximum Gaussian kernel to reduce the impact of inter-subject variability in brain anatomy. Note that all ROI-to-ROI analyses were based on unsmoothed data; only ROI-to voxel seed-based analyses utilized smoothed data.

To address the spurious correlations in resting-state networks caused by head motion, we identified problematic time points during the scan using Artifact Detection Tools (ART, www.nitrc.org/projects/artifact_detect/). Images were specified as outliers based on a criterion of framewise displacement > .4 mm or global signal intensity > 4 standard deviations above the mean signal for that session. Outlier images were not deleted from the time series, but rather, modeled in the first level general linear model (GLM) in order to keep intact the temporal structure of the data. Each outlier was represented by a single regressor in the GLM, with a 1 for the outlier time point and 0 elsewhere. Participants were removed from additional analysis steps if they had greater than 10% of timepoints tagged as outliers (three were removed based on excessive motion).

Using CONN software (Whitfield-Gabrieli & Nieto-Castanon, 2012), physiological and other spurious sources of noise were estimated and regressed out using the anatomical CompCor method (Behzadi, Restom, Liau, & Liu, 2007). Global signal regression was not used due to fact that it mathematically introduces negative correlations (Murphy et al., 2009). The normalized anatomical image for each participant was segmented into white matter (WM), gray matter, and CSF masks using SPM8. To minimize partial voluming with gray matter, the WM and CSF masks were eroded by one voxel. The eroded WM and CSF masks were then used as noise ROIs. Signals from the WM and CSF noise ROIs were extracted from the unsmoothed functional volumes to avoid additional risk of contaminating WM and CSF signals with gray matter signals. The following nuisance variables were regressed out: three principal components of the signals from the WM and CSF noise ROIs; head motion parameters (three rotation and three translation parameters) along with their first-order temporal derivatives; each artifact outlier image; linear trends; and the main effect of each task session and its first-order temporal derivative, which models the initial ramping up of activation the beginning of each session. A band-pass filter (0.009 Hz < *f* < 0.10 Hz) was simultaneously applied to the BOLD time series during this step.

### Network nodes

We used a set of 400 cortical nodes from a recent parcellation (Schaefer et al., 2017) that combined the parcellation approaches of Yeo et al. (2011) and Gordon et al. (2014). We additionally used a select group of subcortical nodes that are relevant to affective processing: 4 nodes spanning the striatum, based on the parcellation of Buckner and colleagues (Buckner, Krienen, Castellanos, Diaz, & Yeo, 2011), 2 nodes for the left and right amygdala from the Harvard-Oxford atlas, and one node covering the periaqueductal gray (PAG) from (Keuken et al., 2014). For each participant, we extracted the mean timeseries from unsmoothed data for each ROI. The residual timeseries (following nuisance regression) was used to compute condition-specific correlation matrices.

### Graph analysis

All graph metrics were computed using the Brain Connectivity Toolbox (Rubinov & Sporns, 2010) on the weighted graphs (Fisher z-transformed correlation coefficients) of individual participants, that were thresholded to retain only positive edge weights.

### Community structure

Many types of complex systems including the brain exhibit modularity, which refers to the fact that nodes tend to be connected with a select group of other nodes, forming distinct modules (also referred to as communities or networks) that perform different functions (Cole, et al., 2014a; Girvan & Newman, 2002; Newman, 2004; Power et al., 2011; Sporns & Betzel, 2016; Yeo et al., 2011). To examine potential reorganization of community structure as a function of cognitive state, we used a modularity maximization algorithm to estimate community structure separately during acceptance and narrative-evaluation. Specifically, we used the Louvain algorithm (Blondel, Guillaume, Lambiotte, & Lefebvre, 2008) implemented with the brain connectivity toolbox (Rubinov & Sporns, 2010). This algorithm finds partitions optimizing the modularity value, Q, by grouping nodes into non-overlapping networks that maximize intra-modular and minimize inter-modular connections (Newman, 2004). The modularity value for weighted graphs, *Q*^*w*^, is computed as follows:

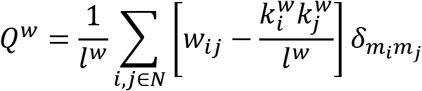

where *w*_*ij*_ is the edge weight between nodes *i* and *j*, *l*^*w*^ is the sum of all weights in the graph, 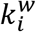 is the weighted degree of node *i*, and *m*_*i*_ is a module containing node i. 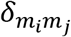 is =1 if nodes *i* and *j* belong to the same module, and = 0 otherwise. The modularity of a partition is a scalar between −1 and 1 that quantifies the strength of within-module edges relative to the strength of between-module edges. The following steps were performed separately for the acceptance and narrative-evaluation data: (i) a group-averaged correlation matrix was generated; (ii) this matrix was submitted to the Louvain algorithm 10,000 times, producing a 1 × 407 vector of community assignments each time; (iii) for each iteration, a 407×407 co-classification matrix was constructed, with entries of 1 or 0 depending on whether nodes *i* and *j* were assigned to the same or different modules; (iv) a final co-classification matrix was generated which reflected the proportion of instances that nodes *i* and *j* were assigned to the same module across the 10,000 iterations; (v) the Louvain community detection algorithm was then applied to this final co-classification matrix to provide a final set of community assignments; (vi) any communities containing fewer than 8 nodes were dissolved, and each node was reassigned to the community to which it showed the strongest mean correlation (outside of the original small community). The number of communities that are detected with this algorithm can be altered with a resolution parameter, gamma (γ), which was set to 2.4, resulting in a 14-network parcellation for each condition. This is comparable to the number of networks that has been observed in prior work (Power, et al., 2011; Yeo, et al., 2011).

### System segregation

We assessed system segregation in two ways. The first, was using clustering. The weighted clustering coefficient, *C*^*w*^, provides a measure of network segregation by quantifying the potential for communication within the immediate neighborhood of a node (Bullmore & Sporns, 2009; Onnela, Saramaki, Kertesz, & Kaski, 2005; Watts & Strogatz, 1998). Specifically, it is defined as the proportion of neighbors around node *i* that are also interconnected (forming a triangle), and is computed as:

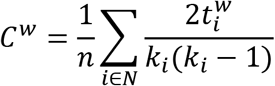

where 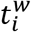 is the number of triangles around node *i*, normalized by their intensity (edge weight), and *k*_*i*_ is the degree (total number of connections) of node *i*. The mean weighted clustering coefficient for the global network is simply the average of all *C*^*w*^values across nodes in the network. Clustering was computed across a range of thresholds (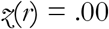 to .99 in .01-step increments) to demonstrate the lack of dependence upon thresholding

Our second measure of system segregation was the segregation index, which reflects the relative strength of within-network to between-network connections (Chan, Park, Savalia, Petersen, & Wig, 2014). It is computed as:

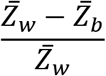

where 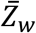 is the mean fisher z-transformed correlation between nodes within the same network, and 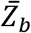is the mean fisher z-transformed correlation between nodes of one system and nodes all other systems. Modular networks display a positive segregation index.

### Global efficiency

Global efficiency, E_glob_, provides a measure of how efficiently information can travel throughout the network, and is defined as the average inverse shortest path distance between each pair of nodes (Achard & Bullmore, 2007; Latora & Marchiori, 2001). Thus, a network with high global efficiency is one in which it is possible to define a path between any pair of nodes with few intervening nodes. It is computed as:

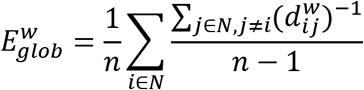

where 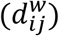 is the shortest weighted path length (distance) between nodes *i* and *j*. In a weighted network, path length is computed as the inverse of weight. Global efficiency was computed across a range of thresholds (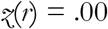 to .99 in .01-step increments) to demonstrate the lack of dependence upon thresholding.

### Hub structure

Hub nodes can be divided into two types: (i) connector hubs, which are central in across-network communication; and (ii) provincial hubs, which are central in within-network communication (Guimera & Amaral, 2005). Connector hubs can be identified with the participation coefficient, which assesses the distribution of a node’s connections across modules. It is computed as:

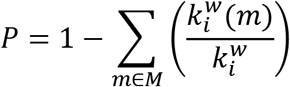

Where *M* is the set of all modules (networks), and 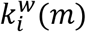 is the strength of connections between node *i* and all nodes in module *m*. A participation coefficient close to 1 indicates that the node’s connections are evenly distributed across all networks, whereas a participation coefficient close to 0 indicates that the node’s connections are confined to its home network.

Provincial hubs can be identified with the within-module degree Z-Score. A node with a high within-module degree Z-score is more connected to members of its own community than other members in that community, thus acting as central node. It is computed as:

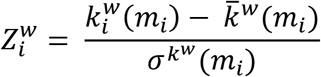

where *m*_*i*_ is the module containing node *i*, 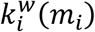 is the weighted within-module degree of *i* (the strength of connections between *i* and all other nodes in *m*_*i*_), and 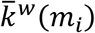 and 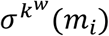 are the mean and standard deviation of weighted degree across nodes in that module.

For each participant and we computed the participation coefficient and within-module degree Z-score for each node across network densities (proportion of retained connections) from .02 to .30 in increments of .01. We then computed the mean value across thresholds. This reduced the possibility that the observed values would be dependent upon a single arbitrarily chosen threshold. We then computed the mean participation coefficient and mean within-module degree Z-score across participants for each node. We then rank ordered nodes to identify those with the highest participation coefficient and highest within-module degree Z-score. We took the 90^th^ percentile and above as connector and provincial hub nodes. If a node was both a connector and a provincial hub, it was considered a connector hub and removed from the collection of provincial nodes, so that the latter exclusively contained nodes that act as local hubs. These analysis steps were performed separately for the narrative-evaluation and acceptance conditions.

### Relationship between functional connectivity patterns and subjective experience

We employed a linear regression approach (Kong et al., 2018) to evaluate the relationship between multivariate FC patterns and acceptance-based introspection scores (i.e., inverse composite narrative-evaluative thinking score during the acceptance condition, reflecting the subjective quality of implementing the acceptance mindset). The guiding inference was that, if a particular functional network is involved in acceptance, then similarities between subjects in functional configuration (i.e., multivariate connectivity weights) of that network should be mirrored by corresponding similarities in acceptance scores. We examined FC patterns in several *a priori* determined networks that were predicted, based on prior research, to mediate processes involved in acceptance-based introspection: the frontoparietal control network (FPCN); the salience network (SN); and the ventral somatomotor network.

We used robust regression (iteratively reweighted least squares with a bisquare weighting function) to predict acceptance scores in individual subjects on the basis of similarity in FC patterns. Robust regression down-weights extreme values and is therefore less influenced by outliers. Following a method similar to Kong et al. (2018), we conducted a regression in which Y values were acceptance scores across subjects, and X values were the sum of acceptance scores for all other subjects, weighted by similarity in FC patterns to the target subject. For example, suppose that *y* is the acceptance score and *p* is the FC pattern of a given target subject. For all other (predictor) subjects, *y*_*i*_ and *p*_*i*_ are the acceptance score and FC pattern for the i’th predictor subject. The regression predicts the target subject’s acceptance score y as:

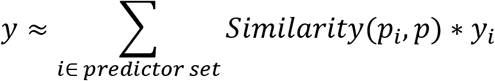

where Similarity(*p*_*i*_, *p*) is the correlation between the FC patterns of the target subject and the i’th predictor subject. FC patterns were the vectorized *multivariate* pattern of all relevant node-to-node FC values (e.g., the vectorized pattern of all node to node FC values within the FPCN). Pearson correlation was used as the similarity metric for FC patterns between each pair of subjects.

This regression model was run separately for each network of interest. A positive beta weight would indicate that subjects with similar acceptance scores also have similar FC patterns. We also examined the relationship between *mean* FC strength within a given network and acceptance scores, which allowed us to determine if the observed relationships were simply due to changes in the strength of FC rather than in the specific pattern of multivariate FC weights. In addition, we included sex and motion (mean framewise displacement) as regressors of no interest in each model. Motion is of particular concern, given the ongoing controversy concerning its ability to result in spurious correlations and temporal variability, with a consequent skewing of brain-behavior relationships (Parkes, Fulcher, Yucel, & Fornito, 2018; Power et al., 2014; Siegel et al., 2017).

### Code availability

Data (connectivity matrices) and code to perform analyses are available at: https://github.com/matthewldixon/Emotional_acceptance.

## RESULTS

### Subjective reports

Participants (*N* = 24) reported different subjective experiences during the two mindsets (**Table 1**). The acceptance condition relative to narrative-evaluative condition was associated with higher scores on awareness of bodily sensations (*t*_23_ = 4.12, *p* < .001), and lower scores on getting caught up in thought, analyzing thoughts/feelings, thinking about the future, and judging thoughts/feelings (paired *t*-test on composite narrative-evaluative thinking score: *t*_23_ = 6.96, *p* < .001). Self-reported log meditation experience was negatively correlated with the composite narrative-evaluative thinking score during the acceptance condition (*r* = −.37, *p* = .038, one-tailed) (**Figure 2A**) and negatively correlated with the extent to which the acceptance mindset was rated as more difficult to implement than the narrative-evaluative mindset (*r* = −.37, *p* = .038, one-tailed) (**Figure 2B).**This potentially suggests that individuals can improve acceptance ability with training. However, our findings cannot differentiate whether individuals with more meditation experience were actually better at implementing the acceptance mindset, or simply believed they were better. Participants exhibiting a greater reduction in narrative-evaluative thinking during acceptance showed a greater increase in awareness of bodily sensations (*r* = −.70, *p* < .001) (**Figure 2C**), suggesting a push-pull relationship between attending to past and future-oriented elaborative thought and judgments, and attending to present-moment bodily sensations. Mood was significantly lower during the narrative-evaluative condition than during a resting state condition, consistent with the instructions to reflect on an upsetting issue (*t*_23_ = 6.55, *p* < .001). Mood was higher during the acceptance condition compared to the narrative-evaluative condition (*t*_23_ = 3.71, *p* = .001) and not statistically different from the resting state condition (*t*_23_ = 1.64, *p* = .11). This could have been due to a habituation effect; however, this seems unlikely given that bodily sensations associated with emotional experience were the strongest during acceptance. Instead, it potentially suggests that mindful acceptance offers a buffering effect against negative mood when confronting an upsetting personal issue. Acceptance was rated as slightly more difficult to implement than the narrative-evaluative mindset, but this difference did not reach significance (*t*_23_ = 1.80, *p* = .069) (**Figure 2D**).

**Table 1.** Post-scan reports

| Variable | Rest | Narrative-evaluative | Acceptance |
| --- | --- | --- | --- |
| Body sensations | 5.13 | 4.28 | 5.70 |
| Mood | 5.26 | 3.78 | 4.70 |
| Caught up in thought |  | 5.02 | 3.85 |
| Analyze situation |  | 4.96 | 2.83 |
| Thoughts of future |  | 4.83 | 2.57 |
| Judge thoughts/feelings |  | 4.09 | 2.30 |
| Difficulty |  | 3.07 | 3.87 |

*Note.* Items were rated on a 7-point likert scale from 1 = not at all, to 7 = a lot / very much (the scale for mood was: 1 = very unpleasant to 7 = very pleasant).

**Figure 2.**
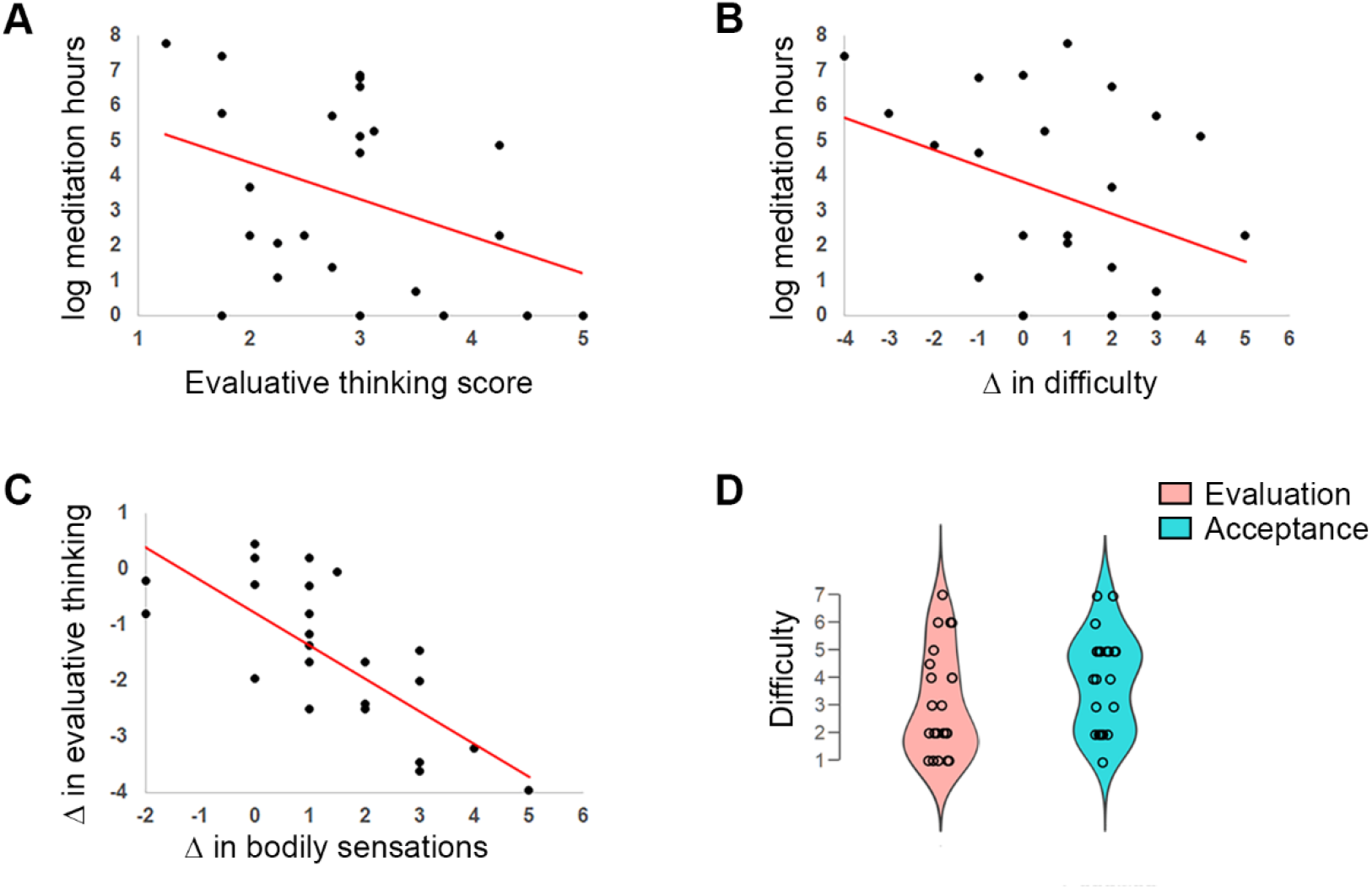
The relationship between narrative-evaluative thinking, meditation experience, bodily sensations, and difficulty. **(A)** Scatter plot showing that a greater number of meditation hours was associated with a lower narrative-evaluative thinking score during acceptance. (**B)** Scatter plot showing that a greater number of meditation hours was associated with less difficulty in implementing the acceptance mindset relative to the narrative-evaluation mindset. **(C)** Scatter plot showing that the greater the decrease in narrative-evaluative thinking from the narrative-evaluation condition to the acceptance condition, the greater the increase in awareness of bodily sensations. (**D**) Violin plots of difficulty scores for narrative-evaluation and acceptance, created with JASP computer software Version 0.9 (JASP Team, 2018).

To place mindful acceptance reports in a broader context, we also had participants complete several questionnaires (see **Supplementary Methods**), and explored the correlations between the measures (**Figure S1**). Descriptively, acceptance scores (inverse of narrative-evaluative thinking scores during the acceptance condition) were positively correlated openness (*r* = .37), hours of meditation experience (*r* = .36), and trait mindfulness (*r* = .26), and negatively correlated with rumination (*r* = −.45), and neuroticism (*r* = −.42). Negative correlations between acceptance reports and rumination (*r* = −.38) and neuroticism (*r* = −.35) were maintained when trait mindfulness scores were partialled out.

### Community structure

To compare brain network organization during the two mindsets, we first used a community detection algorithm to examine how nodes organized into communities. The resulting community structures were similar across conditions (**Figure 3**) and consistent with the major features of prior network parcellations (Power, et al., 2011; Yeo, et al., 2011). However, one notable difference from prior resting state parcellations was that the FPCN fractionated into separate left and right hemisphere networks during both the narrative-evaluative and acceptance mindsets. Notably this pattern was not observed when the community detection algorithm was run on resting state data from the same participants (**Figure S2**). Beyond the general similarity across conditions, there were several striking differences.

**Figure 3.**
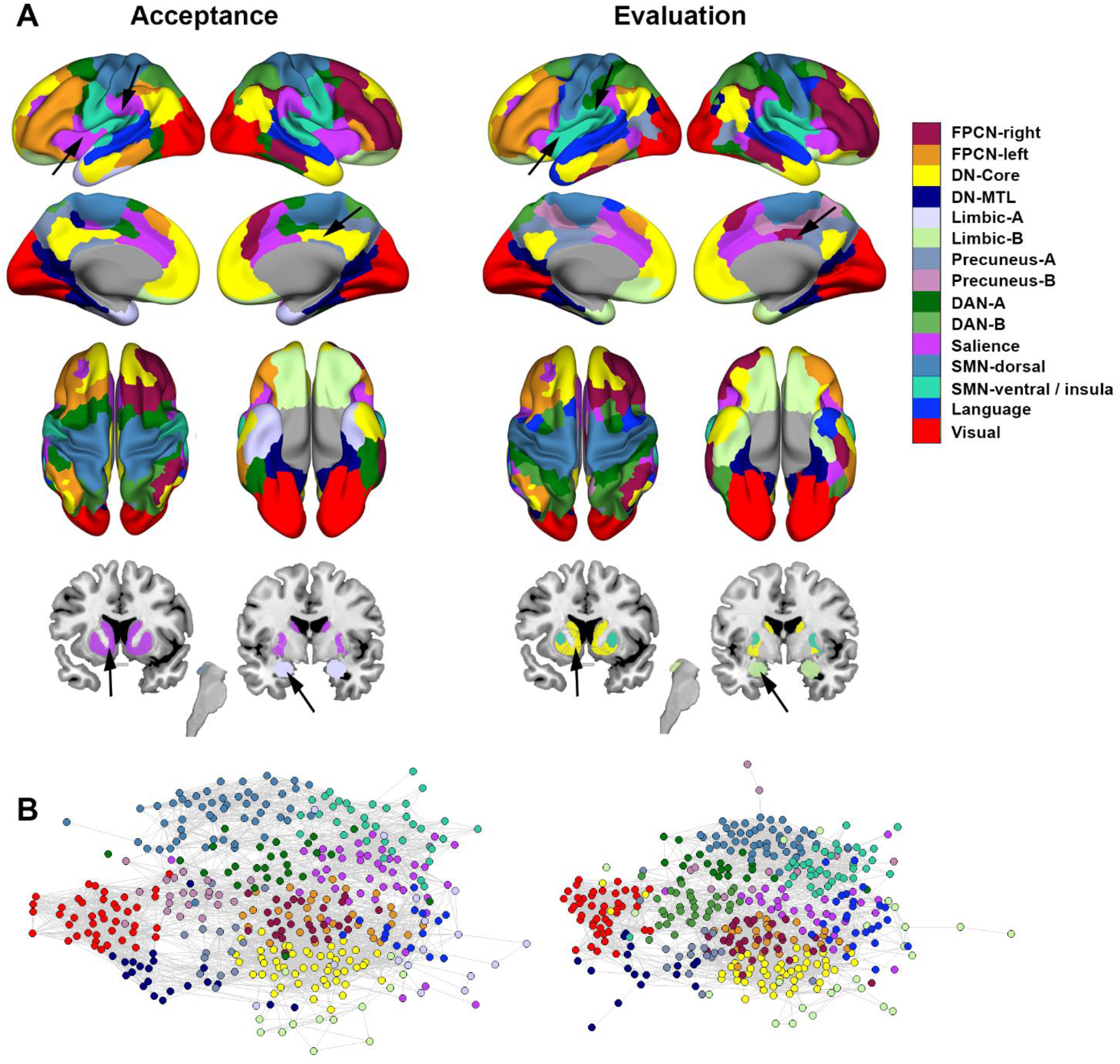
Community structure during the narrative-evaluative and acceptance conditions. **(A)** Surface rendering of the network parcellation based on the community detection algorithm. Arrows highlight points of divergence in community structure across conditions. **(B)** Visualization of the network topology. The group-averaged correlation matrix for each condition (5% density) was submitted to the Kamada–Kawai energy algorithm (Kamada & Kawai, 1989), implemented in Pajek software (De Nooy, Mrvar, & Batagelj, 2011), which produces spring-embedded layouts that minimize the geometric distances of nodes based on their topological distances in the graph. Well-connected nodes are pulled towards each other, whereas weakly-connected nodes are pushed apart in a manner that minimizes the total energy of the system. Nodes are colour-coded based on community assignment. *Abbreviations*: FPCN, frontoparietal control network; DN, default network; MTL, medial temporal lobe subsystem; DAN, dorsal attention network; SMN, somatomotor network.

First, there was a reorganization of interoceptive circuits. During the narrative-evaluative condition, the mid and posterior insula nodes grouped together, forming their own segregated network. In contrast, during mindful acceptance, the mid-insula grouped with the salience network, while the posterior insula grouped with the ventral somatomotor network. Thus, cortical interoceptive regions were more integrated into large-scale systems during acceptance. Additionally, the salience network incorporated more of the rostral supramarginal gyrus near the secondary somatosensory cortex (SII) during acceptance. These changes reveal a state-dependent shift in the organization of large-scale neural circuits underlying the processing visceral and somatosensory information.

Second, there was a reorganization of motivational circuits. The striatum plays a central role in motivation and value-based learning and decision making (Bartra, McGuire, & Kable, 2013; Everitt & Robbins, 2005; Frank, 2005; Haber & Behrens, 2014; Leotti & Delgado, 2014; O’Doherty et al., 2004). During narrative-evaluative processing, the striatum (excluding the dorsal putamen) grouped with the default network, which plays a role in elaborative self-referential thinking and evaluation. In contrast, during acceptance, the striatum grouped with the salience network, which plays a role in interoception and detecting a wide array of currently salient stimuli. To further illuminate this change in network affiliation, we performed a whole-brain seed-based analysis and found that a striatum seed region exhibited more extensive functional coupling with the anterior insula, mid-insula, right inferior frontal sulcus (IFS)/dorsolateral prefrontal cortex (DLPFC), orbitofrontal cortex (OFC), and dopaminergic midbrain (near the substantia nigra/VTA) during acceptance compared to narrative-evaluative processing. On the other hand, the striatum seed region exhibited more extensive coupling with the rostromedial prefrontal cortex (RMPFC) and left IFS during narrative-evaluative processing compared to acceptance (**Figure 4**). This shift in the striatum’s network affiliation may reflect the change across conditions in the motivational relevance of self-referential thinking versus viscerosensory experience. There was also a change in the organization of the limbic network. During the narrative-evaluative mindset, there was a unified limbic network consisting of the orbitofrontal cortex (OFC), amygdala, medial temporopolar cortex, and periaqueductal gray (PAG). In contrast, during acceptance, this network dissolved, with the OFC forming its own distinct network, and the PAG joining the precuneus network.

**Figure 4.**
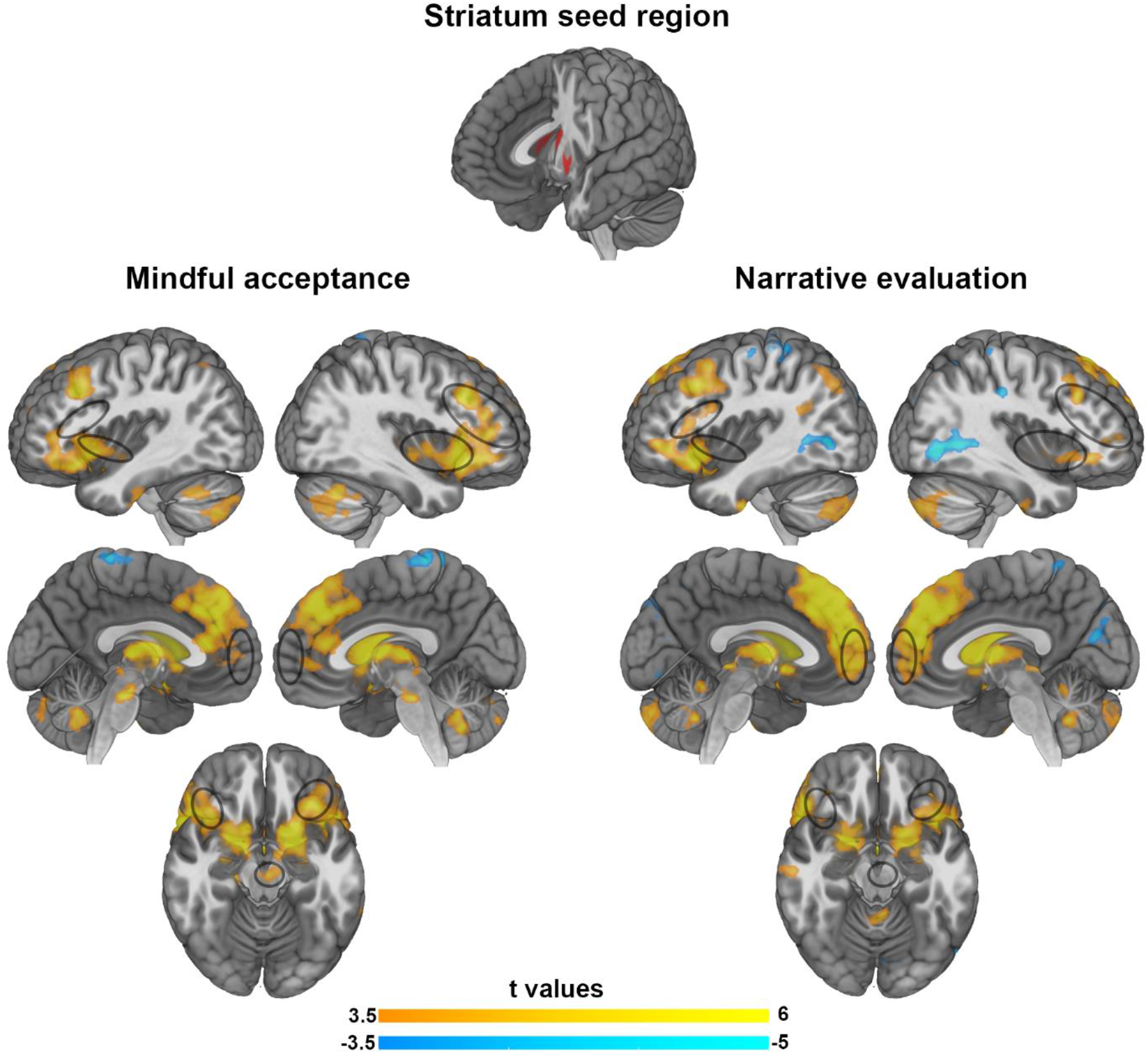
Whole-brain functional connectivity maps using a striatum ROI as the seed. The striatum ROI (which included the caudate body and part of the putamen) showed more extensive functional coupling with the mid insula, right inferior frontal sulcus (IFS)/dorsolateral prefrontal cortex (DLPFC), orbitofrontal cortex (OFC) and dopaminergic midbrain (near the substantia nigra/VTA) during acceptance compared to narrative-evaluative processing. It showed more extensive coupling with the rostromedial prefrontal cortex (RMPFC) and left IFS during narrative-evaluative processing compared to acceptance. Maps were thresholded using a cluster defining threshold of *Z* > 3.1 and a cluster level threshold of *p* < .05, false discovery rate (FDR) corrected. Maps were visualized using MRIcroGL (https://www.nitrc.org/projects/mricrogl/).

Finally, there was a reorganization of the default network. During acceptance, the default network expanded in territory and included the dorsal posterior cingulate cortex (PCC), a broader extent of the posterior inferior parietal lobule (pIPL), and stretched further into the left medial OFC/subgenual anterior cingulate cortex (sgACC). To summarize, there was general stability in the organization of nodes into modular systems, however, there was also evidence of dynamic state-dependent reorganization of nodes involved in interoception, motivation, and self-referential processing.

### System Segregation

While the network parcellation revealed the spatial organization of nodes into communities, additional insights into the network’s capacity for modular processing can be gleaned by quantifying system segregation. Clustering provides a measure of the strength of communication within a tightly interconnected neighborhood of a node, and may provide an index of the capacity for specialized processing within a segregated community. We found no difference between acceptance and the narrative-evaluation condition in the mean strength of clustering (paired *t*-test on mean clustering values derived from unthresholded graphs: *t*_19_ = 1.24, *p* = 0.23) (**Figure 5**). We next examined the segregation index (Chan, et al., 2014), which reflects the relative strength of within-network connections to between-network connections. There was no difference across conditions in the segregation index (paired *t*-test: *t*_19_ < 1) (**Figure 5**).

**Figure 5.**
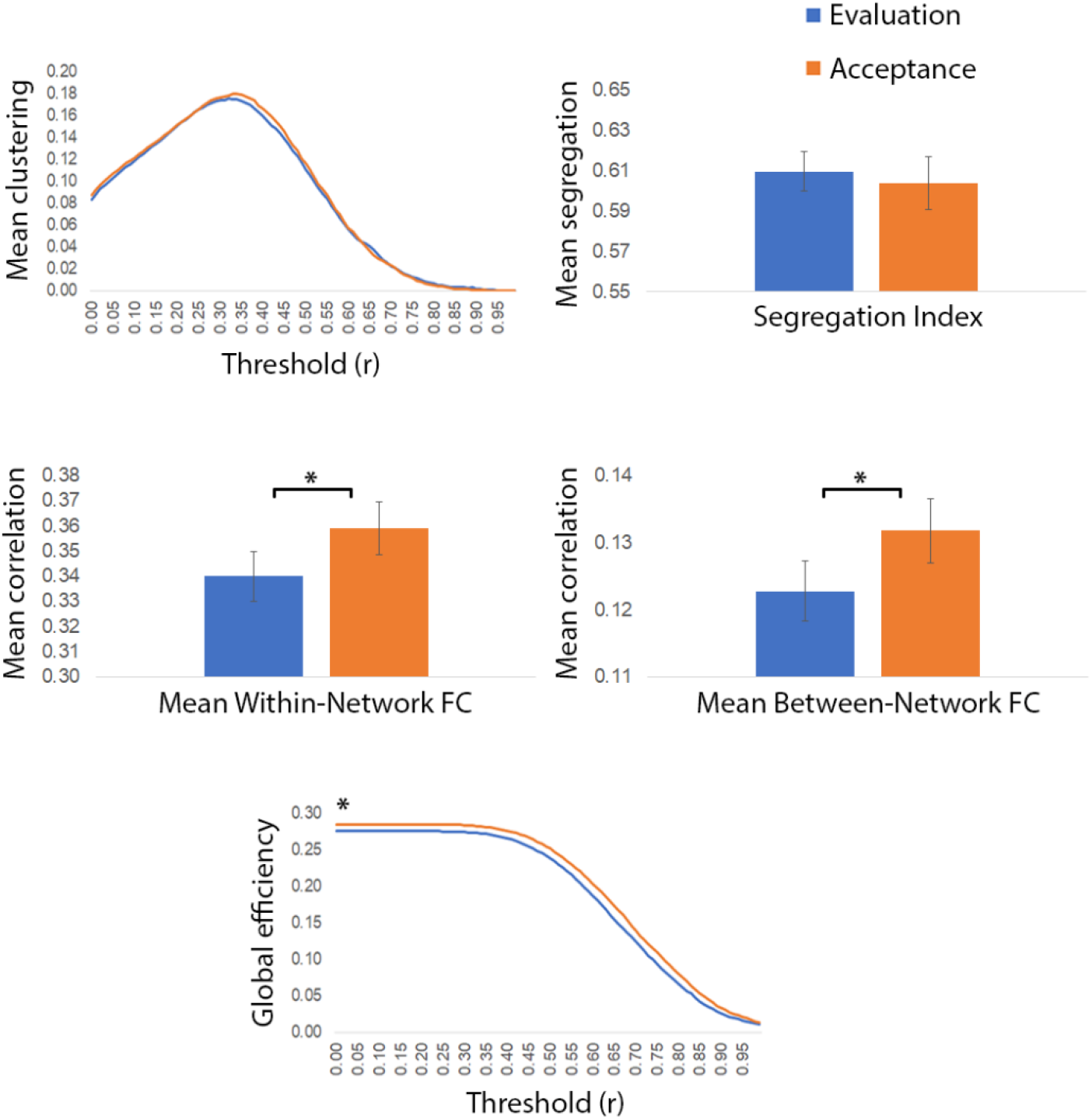
System segregation and global efficiency. There were no differences across conditions in mean clustering or the strength of the segregation index. Mindful acceptance relative to the narrative-evaluative condition was associated with stronger mean within-network and between-network functional connectivity and increased global efficiency, indicating more short pathways between nodes across the network. We examined clustering and global efficiency across a range of thresholds to show the lack of dependence upon thresholding. * denotes *p* < .05.

### Global Connectivity

Global efficiency reflects the number of short pathways between nodes (irrespective of community assignment), and by extension, the ease of information travel across the global network. Acceptance was associated with a significant increase in global efficiency compared to the narrative-evaluative mindset (paired *t*-test on EGlob values derived from unthresholded graphs: *t*_19_ = 2.13, *p* = .047) (**Figure 5**). Consistent with this, acceptance was associated with higher within-network functional connectivity (*t*_19_ = 2.23, *p* = 0.038) and higher between-network functional connectivity (*t*_19_ = 2.17, *p* = 0.043) (**Figure 5)**. A shift towards a more globally interconnected landscape is evident in the group-averaged correlation matrices, which reveal widespread increases in the strength of node-to-node correlations during acceptance versus narrative-evaluation (**Figure 6**).

**Figure 6.**
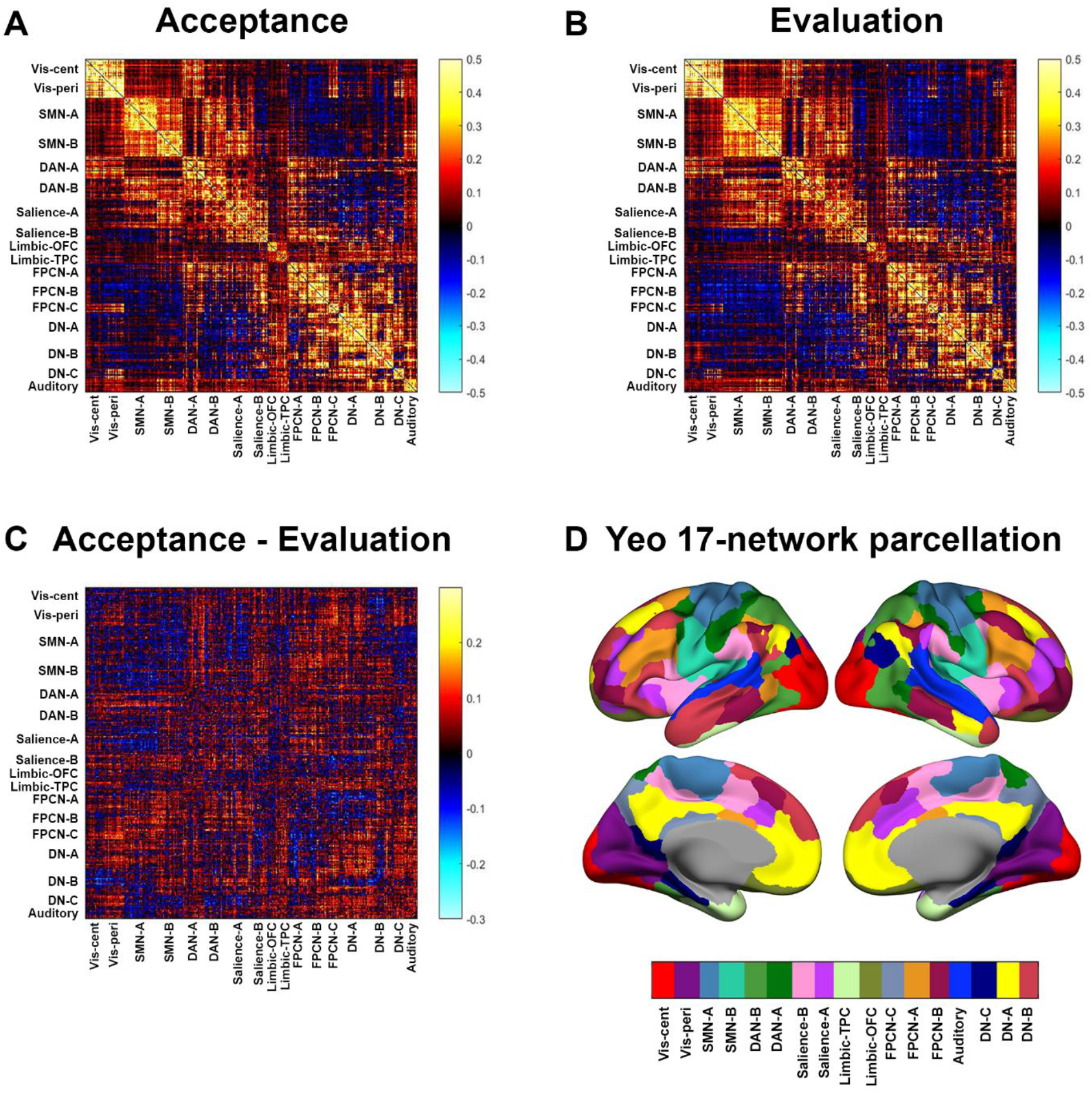
Group-averaged correlation matrices for narrative-evaluation and acceptance. **A**. Group-averaged correlation matrix for acceptance. **B**. Group-averaged correlation matrix for narrative-evaluation. **C**. Group-averaged difference matrix (acceptance minus narrative-evaluation). There is considerably more warm color throughout the difference matrix indicating stronger functional connectivity across much of the global network during acceptance versus narrative-evaluation. **D**. The nodes in each matrix are organized based on the assignments of the 400 cortical nodes to the Yeo 17-network parcellation (Schaefer, et al., 2017). Using an independent parcellation made it possible to organize nodes in the narrative-evaluation and acceptance matrices in an identical manner, thus providing a matched comparison of functional connectivity values during these conditions. Color bar indicates Fisher r-to-z transformed correlation values.

### Hub structure

Networks often contain central ‘hub’ nodes that coordinate information flow across spatially distributed regions (Bertolero, Yeo, & D’Esposito, 2017; Cole, et al., 2013; Hagmann et al., 2008; Power, Schlaggar, Lessov-Schlaggar, & Petersen, 2013; van den Heuvel & Sporns, 2011, 2013). Connector hubs have a high participation coefficient indicating that their connections are widely distributed across multiple brain networks. Connector hub regions that were common to both conditions were mainly located in the frontoparietal control network (FPCN) and dorsal attention network (DAN), consistent with the spatial distribution of hubs observed in prior work (Bertolero, et al., 2017; Gordon et al., 2018; Power, et al., 2013) (**Figure 7**).

**Figure 7.**
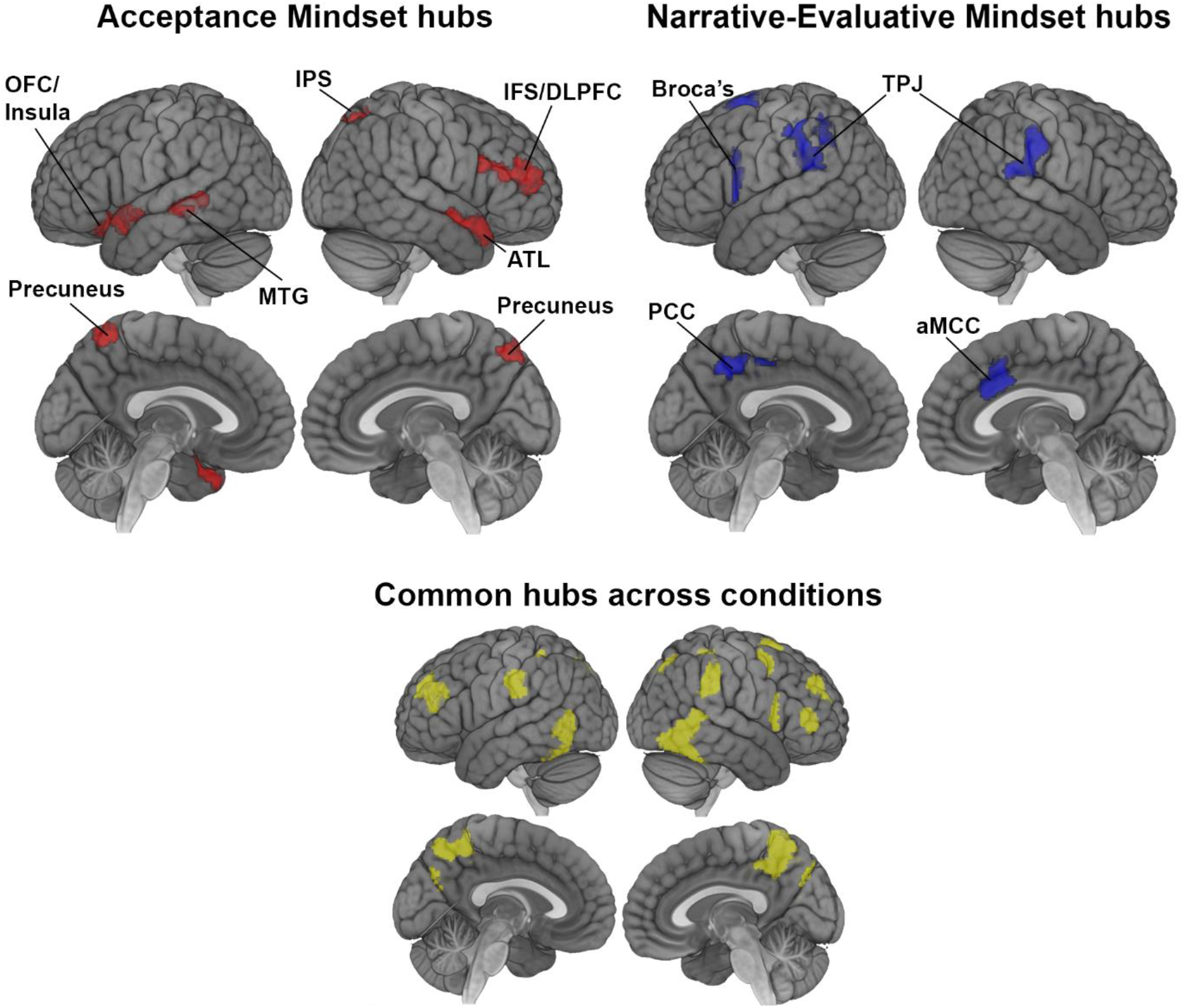
Nodes identified as connector hubs based on their participation coefficients. Nodes highlighted in red were identified as connector hubs during mindful acceptance but not during narrative-evaluative processing. Nodes highlighted in blue were identified as connector hubs during narrative-evaluative processing but not during mindful acceptance. Nodes highlighted in yellow were identified as connector hubs during both conditions. *Abbreviations*: IFS/DLPFC, inferior frontal sulcus/dorsolateral prefrontal cortex; IPS, intraparietal sulcus; PCC, posterior cingulate cortex; OFC, orbitofrontal cortex; MTG, middle temporal gyrus; ATL, anterior temporal lobe; aMCC, anterior mid-cingulate cortex; TPJ, temporoparietal junction

There were also key differences across conditions (**Figure 7**). Connector hubs that were identified during the narrative-evaluative condition but not during the acceptance condition included parts of the default network (left posterior cingulate cortex, bilateral temporoparietal junction), as well as the left inferior frontal gyrus near Broca’s area, and the anterior mid-cingulate cortex. These regions are involved in self-referential processing, mentalizing, and language/conceptual processing. On the other hand, connector hubs that were identified during acceptance but not narrative-evaluation included the regions of the frontoparietal control network that are involved in cognitive control and metacognitive awareness including the right inferior frontal sulcus (IFS), right mid-dorsolateral prefrontal cortex (mid-DLPFC), right intraparietal sulcus (IPS), bilateral precuneus, and left middle temporal gyrus (MTG). Other connector hubs included the left lateral OFC/anterior insula and bilateral anterior temporal lobe (ATL).

Provincial hubs have a high within-module degree Z-score, indicating that they have stronger connections within their home network than other nodes in that network. Provincial hubs common to both conditions included default network regions (medial prefrontal cortex, left posterior inferior parietal lobule), posterior insula, and parts of the visual and somatomotor networks (**Figure S3**). Provincial hubs that were identified during narrative-evaluation but not acceptance included a small area of the right dorsal anterior insula, the left lateral prefrontal cortex (IFS/mid-DLPFC), and parts of visual cortex (**Figure S3**). Provincial hubs that were identified during acceptance but not narrative-evaluation included the left rostrolateral prefrontal cortex, the bilateral anterior and mid insula, bilateral posterior cingulate cortex, and left retrosplenial cortex/calcarine sulcus (**Figure S3**). Thus, while the posterior cingulate cortex was a connector hub linking multiple different networks during narrative-evaluation, it played a more limited role as a central node within its home network (the default network) during acceptance. Additionally, the mid insula became a central node within the salience network during acceptance but not during evaluation-narrative processing. These findings reveal a widespread reorganization of hub structure based on cognitive state during emotional processing.

### Relationship between functional connectivity patterns and subjective experience

We next conducted an exploratory analysis to look for a relationship between subjective reports of acceptance and FC patterns in several networks of interest. Using robust regression, we found a statistically significant positive relationship between multivariate FC patterns within the left frontoparietal control network (FPCN) and acceptance scores: subjects demonstrating greater similarity in acceptance scores also demonstrated greater similarity in FC values across nodes within the FPCN (*β* = 0.48, *p* = 0.03; **Figure 8**). Conversely, subjects demonstrating greater similarity in acceptance scores demonstrated lower similarity in FC values across nodes within the salience network: (*β* = 0.49, *p* = 0.04). No relationship was found between acceptance scores and FC patterns within the other networks of interest [right FPCN: (*β* = 0.1, *p* = 0.64); ventral somatomotor network: (*β* = −0.36, *p* = 0.17)] or with FC patterns between networks (all *p*’s > .08).

**Figure 8.**
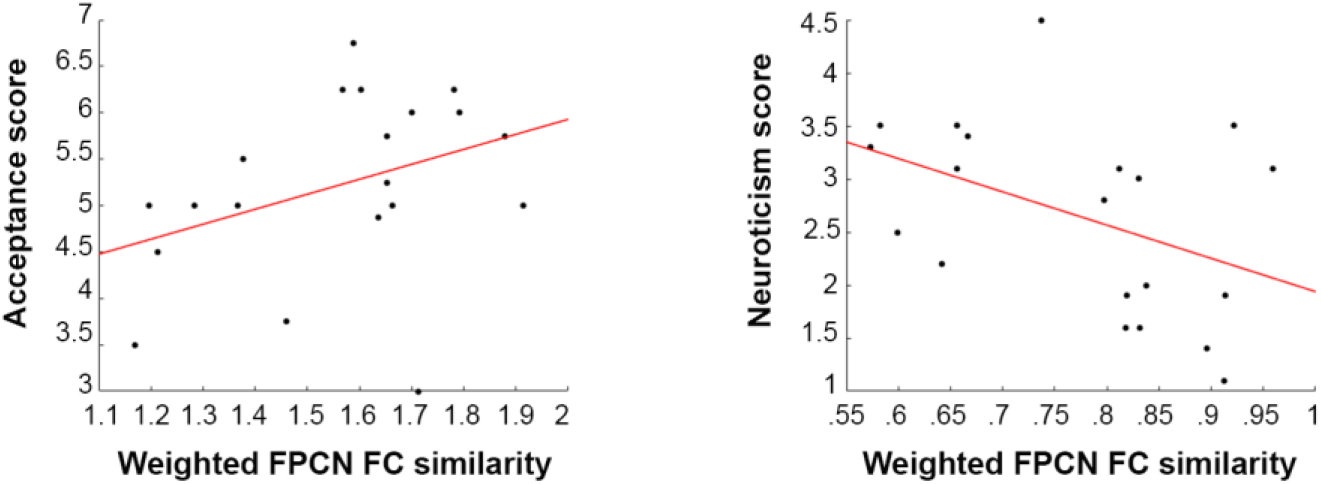
Scatter plots displaying the relationship between the left frontoparietal control network (FPCN) weighted similarity scores and acceptance scores (left panel) and neuroticism scores (right panel). The acceptance score of the target subject (Y axis) was predicted by the sum of acceptance scores for all other subjects weighted by the similarity of FPCN functional connectivity patterns between each predictor subject and the target subject (X axis). A positive relationship indicates that subjects with more similar acceptance scores also had more similar FPCN FC patterns. On the other hand, the negative relationship with neuroticism indicates that subjects with more similar neuroticism scores had less similar FPCN FC patterns.

Given that the left FPCN exhibited the expected positive relationship with acceptance, we further explored the functional relevance of FC patterns within this network. Specifically, we looked for relationships with neuroticism and rumination – traits that are counter to mindful acceptance and are characterized by the tendency to engage in compulsive evaluative thinking in response to actual or anticipated negative experiences (Cattell & Scheier, 1961; Nolen-Hoeksema, 2000). Accordingly, we expected to find an inverse relationship between left FPCN patterns and neuroticism and rumination tendency. Consistent with this, we found a statistically significant negative relationship between multivariate FC patterns within the left FPCN and neuroticism scores: subjects demonstrating greater similarity in neuroticism scores demonstrated less similarity in FC values across nodes within the FPCN (*β* = −0.40, *p* = 0.02; **Figure 8**). There was no relationship between FPCN FC patterns and rumination scores (*β* = −0.06, *p* =0.81). Finally, we found no relationship FPCN FC patterns and trait mindfulness scores (*β* = 0.03, *p* = 0.90), suggesting that FPCN FC patterns during the acceptance condition were specifically related to subjective reports of acceptance quality during the task. Notably, there was no relationship between the mean strength of FC within the left FPCN during the acceptance condition and acceptance scores (*β* = −0.02, *p* = 0.94). This suggests that important information is contained within the specific patterns of interregional connectivity of the left FPCN and not in overall strength of connections. However, these results should be interpreted with caution in light of our modest sample size.

### Relationship to task difficulty

Participants generally found acceptance to be the harder mindset to implement. Thus, it remains possible that some findings were driven by difficulty rather than the acceptance cognitive state. In particular, the global increase in functional connectivity and the shift towards frontoparietal regions as hubs could have reflected additional effort or arousal instead of a qualitative shift in mindset. To examine this issue, we correlated subjective reports of difficulty with these measures. Difficulty difference scores (acceptance difficulty - narrative-evaluative difficulty) were not associated with global efficiency values (*r* = −.01, *p* = .97), but were negatively correlated with the mean participation coefficient across frontoparietal control network (FPCN) nodes (left FPCN: *r* = −.56, *p* = .01; right FPCN: *r* = −.48, *p* = .03). Similar findings were obtained when using absolute difficulty ratings during acceptance (global efficiency: *r* = −.19, *p* = .42; mean participation coefficient for left FPCN: *r* = −.47, *p* = .037; mean participation coefficient for right FPCN: *r* = −.36, *p* = .12). These findings are limited by our small sample size, but if anything, suggest that increased global connectivity and a shift towards frontoparietal hubs is associated with *lower* difficulty during acceptance (and perhaps more effective implementation of the acceptance mindset).

### Relationship to meditation experience

An interesting question is whether individuals with more meditation experience (and possibly more experiencing adopting an acceptance-based mindset) require less FPCN involvement given that acceptance may become more of a natural mindset during emotional experience. Alternatively, FPCN involvement may be a key aspect of implementing the acceptance mindset irrespective of training. In line with the latter notion, we did not observe a significant relationship between log meditation hours and the mean participation coefficient across FPCN nodes during acceptance (left FPCN: *r* = .27, *p* = .25; right FPCN: *r* = .19, *p* = .42). This finding is limited by our small sample size and limited variability in meditation experience across participants, but tentatively suggests that meditation experience may not be associated with a reduction in the involvement of FPCN regions during acceptance.

## Discussion

Mindful acceptance and narrative-evaluative processing are two fundamentally different strategies for coping with challenging emotions. Acceptance involves a broadening of attentional scope such that a continuous flow of diverse sensory signals is registered in consciousness. In contrast, narrative-evaluation involves a narrowing of attentional scope around a limited band of sensory signals, with consciousness being predominantly occupied with cognitive elaborations and judgments related to the causes and consequences of one’s feelings. The results of the current study provide evidence that specific brain network properties reorganize to support these distinct mindsets. Specifically, we found that acceptance compared to narrative-evaluation was associated with: (i) an increase in global network connectivity; (ii) a reorganization of community structure, with interoceptive regions becoming more integrated into large-scale networks; (iii) a shift in hub regions, with default network and language regions becoming less central, and FPCN regions becoming more central; and (iv) a shift in motivational circuits, including a change in the striatum’s affiliation from the default network to the salience network, and the dissolution of a limbic network involving the amygdala and orbitofrontal cortex. We also found a relationship between multivariate patterns of FC weights across nodes within the left FPCN and subjective reports of acceptance. These results provide insight into the network dynamics that underly two distinct mindsets that have implications for emotional health and the treatment of affective disorders (Campbell-Sills, et al., 2006; Farb, Anderson, Bloch, & Segal, 2011; Goldin, et al., 2013; Hayes, et al., 2006; Hofmann, et al., 2010; Nolen-Hoeksema, 1991; Segal, et al., 2002).

### Task-dependent network reconfiguration

A large amount of research has been aimed at delineating the functional network organization of the human brain (e.g., Power et al., 2011; Yeo et al., 2011). This work has, by and large, been based on analyzing brain activity during the ‘resting state’, guided by the assumption that this state, devoid of a particular task, is best suited to reveal intrinsic, task-independent brain network organization (Raichle & Snyder, 2007; Smith et al., 2009; Sporns, 2014). Recent work, however, has challenged the notion of ‘intrinsic’ connectivity, instead arguing that the ‘resting state’ is not a baseline state of brain operation, but simply another mental state with its own idiosyncratic cognitive phenomena such as mind wandering and memory recall (Buckner et al., 2013; Krienen et al., 2014). Indeed, a large number of studies have now systematically documented changes in brain network properties as a function of time and task state (Fornito, et al., 2012; Gonzalez-Castillo & Bandettini, 2017; Hutchison, et al., 2013; Medaglia, et al., 2015; Metzak, et al., 2011; Shine, et al., 2016). In the present study, we add to this literature, and demonstrate marked differences in the organization of brain networks as a function of the mindset applied to emotional experience.

### Increased global connectivity during acceptance

We found that acceptance relative to narrative-evaluation was associated with increased global efficiency, an index of the ease of information flow across the global network. Further inspection revealed that acceptance was associated with stronger within-network connections and stronger between-network connections. A recent study found a global increase in between-network interactions when individuals used cognitive reappraisal to down-modulate negative emotion (Brandl, et al., 2018). Other studies have found increased global efficiency (Cohen & D’Esposito, 2016) and increased global network integration (Shine, et al., 2016) during an N-back working memory task relative to rest. These findings suggest that enhanced global communication may be a common feature of network organization during tasks that require cognitive control. A more globally interconnected state during acceptance may enable a broad, reflective monitoring of present moment sensory experience. Additionally, this architecture may reflect the cognitive control operations required to override the tendency of avoiding uncomfortable feelings and getting caught up in elaborative thinking and judgments – ostensibly the default mode of introspection during the experience of challenging emotions.

Another finding that is consistent with a broadened attentional scope during acceptance was the shift in the organization of the default network. The data-driven network parcellation revealed that the default network encompassed the dorsal PCC during acceptance but not during narrative-evaluation. The connectivity patterns and functions of the dorsal PCC contrast with those of the ventral PCC (Leech & Sharp, 2013). The dorsal PCC exhibits connectivity with default, somatomotor, and frontoparietal control networks (Leech, Braga, & Sharp, 2012). This suggests that the default network may receive a greater influx of input from diverse networks during acceptance via the dorsal PCC. The dorsal PCC may regulate the balance of internal and external attention, and may be particularly engaged when individuals adopt a broad (rather than narrow) internal focus (Leech 2013). Thus, by joining the default network during acceptance, the dorsal PCC may shift the dynamics in this network to promote a mode of internal attention characterized by a broad monitoring of inputs rather than a narrow focus on a specific stream of thought related to the meaning and desirability of one’s feelings. Notably, although the territory (spatial extent) of the default network was larger during acceptance compared to narrative-evaluation, this may occur in the context of reduced activation levels and less influence overall, given prior findings of a relative deactivation of the default network during acceptance and meditative states (Brewer, et al., 2011; Farb, et al., 2007; Vago & Zeidan, 2016).

### Reorganization of interoceptive circuits

Acceptance and narrative-evaluation fundamentally differ in the nature of interoceptive processing. Whereas acceptance emphasizes the experience of ‘raw’ sensory signals from within the body as they change from moment-to-moment, narrative-evaluation emphasizes the elaborative meaning and value of those sensations. A key finding revealed in our data-driven network parcellation was a reorganization of interoceptive circuits as a function of mindset. The mid and posterior insula have been identified as primary interoceptive cortex (Craig, 2002; Farb, et al., 2012b) and represent bodily signals (e.g., respiratory and cardiac sensations) based on afferent inputs from homeostatic control regions including the periaqueductal gray, parabrachial nucleus, and ventromedial thalamus (Craig, 2002; Critchley & Harrison, 2013; Damasio & Carvalho, 2013; Farb, et al., 2012b). During narrative-evaluation, these regions formed their own network and were segregated from other large-scale networks. Less integration with other networks suggests a relative compartmentalization of bodily sensations and perhaps exclusion from awareness. With attention directed to cognitive elaborations and meaning making, signals from these insular subregions may not propagate to other nodes, lessening their influence and access to consciousness. This is consistent with participants’ subjective reports which revealed an opposing relationship between attending to bodily sensations versus narrative-evaluative thought. An interesting consequence is that high-level appraisals and the generation of meaning may occur in a context that is relatively devoid of the concrete sensory signals that initiated the cognitive elaborations in the first place.

In contrast, these insular subregions were integrated into large-scale networks during acceptance. Specifically, the mid insula was incorporated into the salience network and the posterior insula was incorporated into the ventral somatomotor network. Greater integration of these insular subregions into large-scale networks during acceptance may allow concrete bodily signals to propagate across distributed neural populations and into consciousness. Indeed, the mid insula became a provincial hub during acceptance, suggesting a central role in salience network information processing. Interestingly, prior studies have reported changes in insula activation patterns and structural integrity following meditation training (Farb, et al., 2010; Farb, et al., 2013; Fox et al., 2016; Fox, et al., 2014; Young, et al., 2017) suggesting that the interoceptive functions of this region may be a key neural substrate underlying present moment awareness of sensory signals.

### Reorganization of motivational circuits

Changes in motivational circuits may also contribute to changes in attention that underlie the instantiation of a narrative-evaluative or acceptance-based mindset. Our network parcellation revealed state-dependent changes in striatum’s network assignment. The striatum, which includes the caudate, putamen, and nucleus accumbens, plays a central role in tagging stimuli with incentive salience during learning and generating the motivational force underlying behavior (Bartra, et al., 2013; Everitt & Robbins, 2005; Haber & Behrens, 2014; Leotti & Delgado, 2014; O’Doherty, et al., 2004). Prior work has shown that striatal functional coupling patterns are flexible (Gerraty et al., 2018) and that striatal reinforcement learning signals influence an integrated priority map that guides attention (Goldfarb, Chun, & Phelps, 2016). Here, we found that the striatum joined the default network during narrative-evaluation, but instead joined the salience network during acceptance. Stronger communication with default network regions including the medial prefrontal cortex may provide a motivational boost that directs attention to elaborate self-referential thoughts and judgments, whereas stronger communication with salience network regions including the insula may provide a motivational boost that directs attention to concrete viscerosensory experience. The coupling patterns of the striatum may therefore influence the nature and scope of attention during emotional experience and flexibility in these patterns may be a key neural substrate by which an acceptance mindset is instantiated.

There was also a state-dependent shift in the organization of the limbic network. The network parcellation delineated a unified limbic network consisting of the orbitofrontal cortex (OFC), amygdala, medial temporopolar cortex (mTPC), and periaqueductal gray (PAG) during narrative-evaluation. Interestingly, this network dissolved during acceptance, with the OFC and amygdala/mTPC forming separate networks and the PAG joining the precuneus network. While the functional significance of this limbic network dissolution is unclear, it reinforces the notion that mindful acceptance and narrative-evaluative processing differ in the organization of motivational circuits.

### Shift from default network to frontoparietal control network regions as hubs

The extent to which attention is narrow or broad in scope may depend on the hub structure of the global brain network. Prior work has uncovered topologically central ‘hub’ regions that share connections with many other nodes, thus providing an infrastructure for the coordination and integration of information across spatially distributed specialized processing modules (Bertolero, et al., 2017; Hagmann, et al., 2008; Power, et al., 2013; van den Heuvel & Sporns, 2011, 2013). Hub regions are prominent in frontoparietal, default, and salience network regions (Baker et al., 2015; Hagmann, et al., 2008; Power, et al., 2013; van den Heuvel & Sporns, 2011) and play a key role in optimizing processing in specialized systems in accordance with task demands (Cole, et al., 2013). During narrative-evaluation but not acceptance, the left posterior cingulate cortex (PCC), bilateral temporoparietal junction (TPJ), Broca’s area, and the anterior mid-cingulate cortex (aMCC) were connector hubs, suggesting considerable influence on network dynamics. The PCC and TPJ are canonical default network regions (Andrews-Hanna, Reidler, Sepulcre, Poulin, & Buckner, 2010) and are activated during autobiographical memory (Andrews-Hanna, et al., 2014a), judgements about personal traits (Denny, Kober, Wager, & Ochsner, 2012), mentalizing (Van Overwalle & Baetens, 2009), and narrative-based self-reflection (Farb, et al., 2007). These region are also associated with getting ‘stuck’ in one’s thoughts (Garrison et al., 2013a) and mind wandering (Christoff, et al., 2009a; Hasenkamp, Wilson-Mendenhall, Duncan, & Barsalou, 2012). During narrative-evaluation, these regions may coordinate information flow across networks such that emotional feelings are colored with elaborative autobiographical details and become incorporated into a temporally-extended narrative, as individuals search for meaning. Additionally, Broca’s area may contribute to the linguistic analysis of internal experience, while the aMCC may contribute to the preparation of defensive emotional behaviors in response to anxiety-provoking thoughts (Shackman et al., 2011; Vogt, 2005, 2009).

The hub structure of the brain exhibited considerable changes during acceptance, as regions of the FPCN became connector hubs. These regions included the mid-dorsolateral prefrontal cortex (mid-DLPFC), inferior frontal sulcus (IFS), intraparietal sulcus (IPS), middle temporal gyrus (MTG), and precuneus. These regions are involved in cognitive control-related processes including working memory and task switching (Cole, et al., 2014b; Dixon, et al., 2018; Dosenbach et al., 2007; Vincent, Kahn, Snyder, Raichle, & Buckner, 2008) and may facilitate a shift away from the default tendency of meaning-making and judgment when confronting challenging emotional feelings. In particular, given the role of mid-DLPFC in protecting task-relevant information in working memory from interference (Sakai, Rowe, & Passingham, 2002), this region may play a key role in maintaining attention on bodily sensations and away from elaborative thoughts during acceptance, thus freeing interoceptive processing from habitual meaning-based interpretations and patterns of reactivity. Moreover, these regions along with the rostrolateral prefrontal cortex – which was a provincial hub during acceptance but not during narrative-evaluative processing – have been linked to metacognitive awareness (Baird, Smallwood, Gorgolewski, & Margulies, 2013; De Martino, Fleming, Garrett, & Dolan, 2013; Fleming, et al., 2010; McCaig, et al., 2011; McCurdy et al., 2013; Ye, Zou, Lau, Hu, & Kwok, 2018) and exhibit altered structure in meditation practitioners (Fox, et al., 2014). The fact that these regions have a strong influence on overall network dynamics during acceptance is consistent with the emphasis on monitoring feelings and sensations with an objective and non-judgmental metacognitive perspective. Thus, a shift towards frontoparietal control network regions as hubs during acceptance may contribute to the ability to flexibly register and then let go of arising mental content (Vago & Zeidan, 2016) and buffer against maladaptive thinking that often arise during the experience of challenging emotions.

### Relationship between functional connectivity patterns in the FPCN and subjective reports

Using a recently developed regression analysis approach for investigating brain-behavior relationships (Kong et al. 2018), we found that multivariate patterns of FC weights across nodes within the left FPCN were associated with subjective reports of acceptance. Specifically, participants showing greater similarity in left FPCN FC patterns also demonstrated greater similarity in acceptance reports. This finding is consistent with the shift in hub structure and indicates a central role for the FPCN in the cognitive mechanisms underlying acceptance. The FPCN is consistently recruited during emotion regulation tasks (Buhle et al., 2014; Dixon, et al., 2020; Kohn et al., 2014; Ochsner & Gross, 2005) and is a primary locus of cognition-motivation interactions (Dixon & Christoff, 2012; Parro, Dixon, & Christoff, 2018; Pessoa, 2008). Of note, a recent study found that the strength of FC between the FPCN and the rest of the brain was negatively correlated with depressive symptoms in non-clinical individuals, which supports a critical role for the FPCN in maintaining mental health and well-being (Schultz et al., 2018).

The fact that the left FPCN in particular demonstrated a relationship with acceptance is interesting in light of prior work linking neural activity in the left PFC to approach-related behaviours typically associated with positive affect (Davidson, Jackson, & Kalin, 2000). Mindful acceptance involves an approach-related orientation towards uncomfortable sensations, with the intent to fully experience whatever arises from moment to moment. One possibility is that the left FPCN contributes to the process of maintaining a consistent approach-related orientation characterized by curious and non-reactive attention to emotional experience. By linking acceptance reports to functional connectivity patterns, this neurophenomenological approach (Lutz & Thompson, 2003) goes a step beyond most previous attempts to link brain to behavior, which have primarily inferred relationships based on lower-level behavioral measures such as reaction time and accuracy during cognitive task conditions.

Interestingly, some prior studies have failed to observe increased FPCN activation during mindful acceptance (Kober, Buhle, Weber, Ochsner, & Wager, 2019; Westbrook et al., 2013), while other work has detected activation (Dixon, et al., 2020; Goldin, et al., 2019). One possibility is that there may be subtle changes in overall signal in the FPCN during mindful acceptance, but more substantial network reorganization – the latter being the primary way that this network makes a contribution to this cognitive state. Indeed, on a theoretical level, the act of intentionally directing attention to emotional experience and exerting effort to disengage from habitual interpretations and reactivity should rely on the functions of the FPCN, given what we know about this network. The findings observed here suggest that changes in the FPCN’s role within the overall network communication structure may be a fruitful target to investigate in future research.

### Limitations

There are several limitations that should be considered when interpreting the results of this study. First, our conclusions are tempered by our modest sample size. While the observed changes in interoceptive, limbic, self-referential, and metacognitive regions make sense in light of the contrast between narrative-evaluation and acceptance, they require validation in future work using a larger sample. Second, our findings may be specific to the characteristics of sample employed here. Although we did not pre-select participants based on meditation experience, many individuals in our study had at least tried meditation, and several individuals had extensive meditation experience. It remains unknown whether similar patterns of network reconfiguration would be observed in a sample consisting entirely of individuals with no meditation experience. A third limitation is that we collected self-reports after scanning rather than online during the conditions which may have led to misremembering and erroneous reports. Additionally, subjective reports can be prone to expectancy and demand effects, and we cannot objectively verify that individuals were able to adopt the distinct modes of introspection (see Van Dam et al., 2017 for further discussion of the limitations of self-reports.). This is a limitation that is difficult to circumvent as these inner experiences do not have clear physiological or behavioral correlates. Physiological and behavioral reactions can be independent of how participants subjectively experience those reactions, thus making it virtually impossible to objectively measure the quality of acceptance. A fourth potential limitation is that participants in our study could have been engaging in different amounts of task-unrelated thinking during the narrative-evaluation and acceptance conditions. Future work employing online experience sampling probes (Christoff, et al., 2009a; Hasenkamp, et al., 2012) could help to rule-out the possibility that differential frequencies of task-unrelated thought contributed to the observed differences in network configuration. A fifth limitation is that we focused on group-level network properties, which do not reveal the full picture, given that there are important individual differences in network properties (Braga & Buckner, 2017; Gordon et al., 2017). A sixth potential limitation is that the acceptance condition always followed the narrative-evaluation condition. We held the order constant because narrative-based meaning making appears to be the more default introspective style of most individuals (Farb, et al., 2007) and we hypothesized that having acceptance come before narrative-evaluation may lead to carryover effects due to the challenge of implementing this type of introspection. It seems unlikely, however, that general condition order effects (e.g., fatigue) could have impacted our results. Fatigue and sleep are associated with reduced strength of within-network connections and less network segregation (Yeo, Tandi, & Chee, 2015). In contrast, we found that acceptance was associated with increased within-network connections and no difference in our measures of segregated processing (clustering and segregation index). Additionally, we observed changes in functional connections that were functionally relevant to narrative-evaluation and acceptance, including greater centrality of frontoparietal regions that are associated with attentional control. A final limitation is that the network basis of acceptance is likely to change across time as a function of experience given that acceptance is a skill that requires training to fully develop. It will therefore be critical for future work to employ longitudinal designs and assess network properties at different stages of expertise with acceptance practice.

### Conclusions

The current study reveals dynamic changes in network topology across narrative-evaluative and acceptance-based emotional processing. Acceptance relative to narrative-evaluation was associated with a more globally interconnected brain network configuration; greater integration of interoceptive insular subregions into large-scale networks; a shift from default network and language regions to frontoparietal control network regions as central hubs that coordinate information flow; and a shift in the organization of motivation circuits. These network changes may underlie the capacity to free interoceptive processing from habitual interpretations, elaborations, and patterns of reactivity, and instead, enable an open mindset characterized by a broad attentional scope and sensory-based experiencing. More broadly, our findings may have implications for understanding the biological basis of clinical disorders such as depression and anxiety that involve maladaptive self-referential processing and heightened emotional reactivity.

## Acknowledgments

We thank Caitlin Mills for generating Figure 1. This work was supported by NSERC (RGPIN 327317-11) and CIHR (MOP-115197) grants to K.C. The sponsors have no involvement in study design; in the collection, analysis and interpretation of data; in the writing of the report; or in the decision to submit the article for publication

