## Supplementary material for "Brain Network Organization During Mindful Acceptance of Emotions"

### Supplementary Information

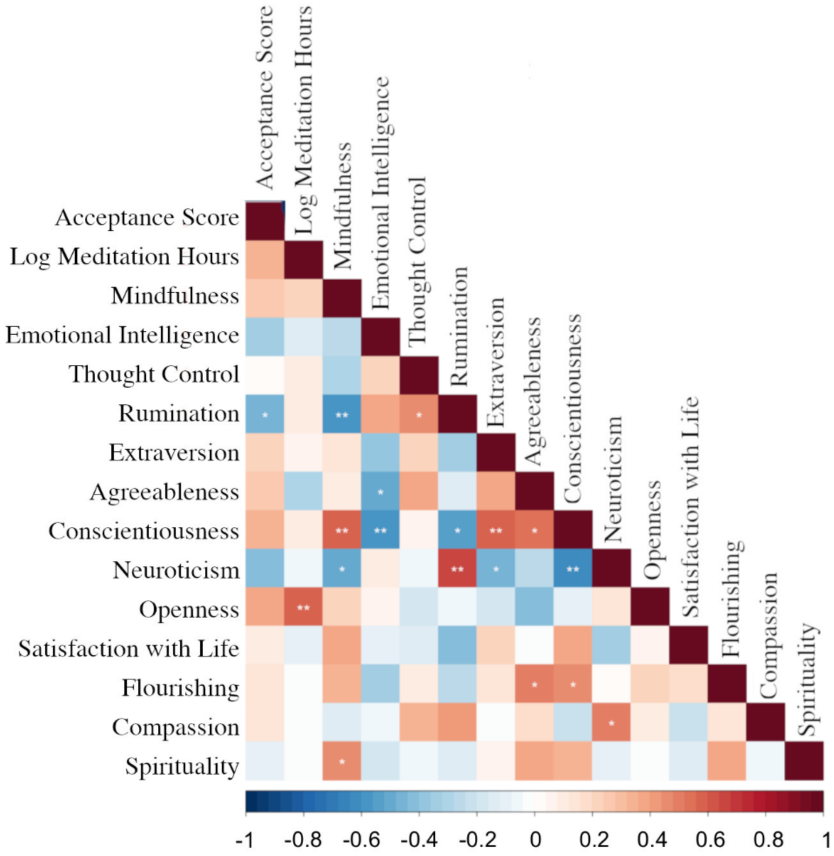


**Figure S1**. Correlations between acceptance reports during the scanning session and responses to post-scanning questionnaires.

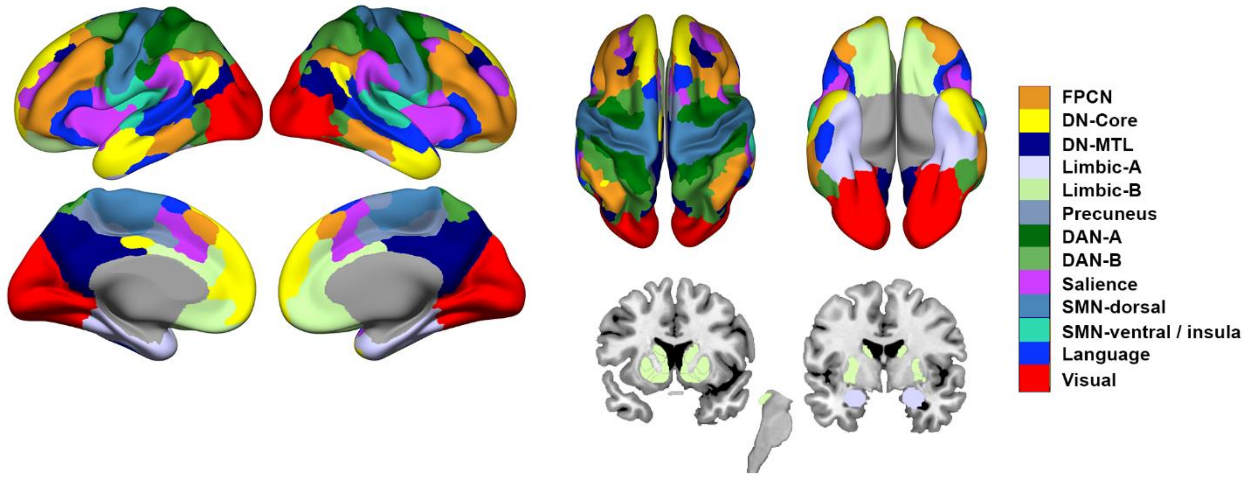


**Figure S2.** Community structure during rest. Surface rendering of the network parcellation based on the community detection algorithm applied to resting state data. In contrast to rumination and acceptance, the FPCN did not fractionate into left and right hemisphere networks during rest.

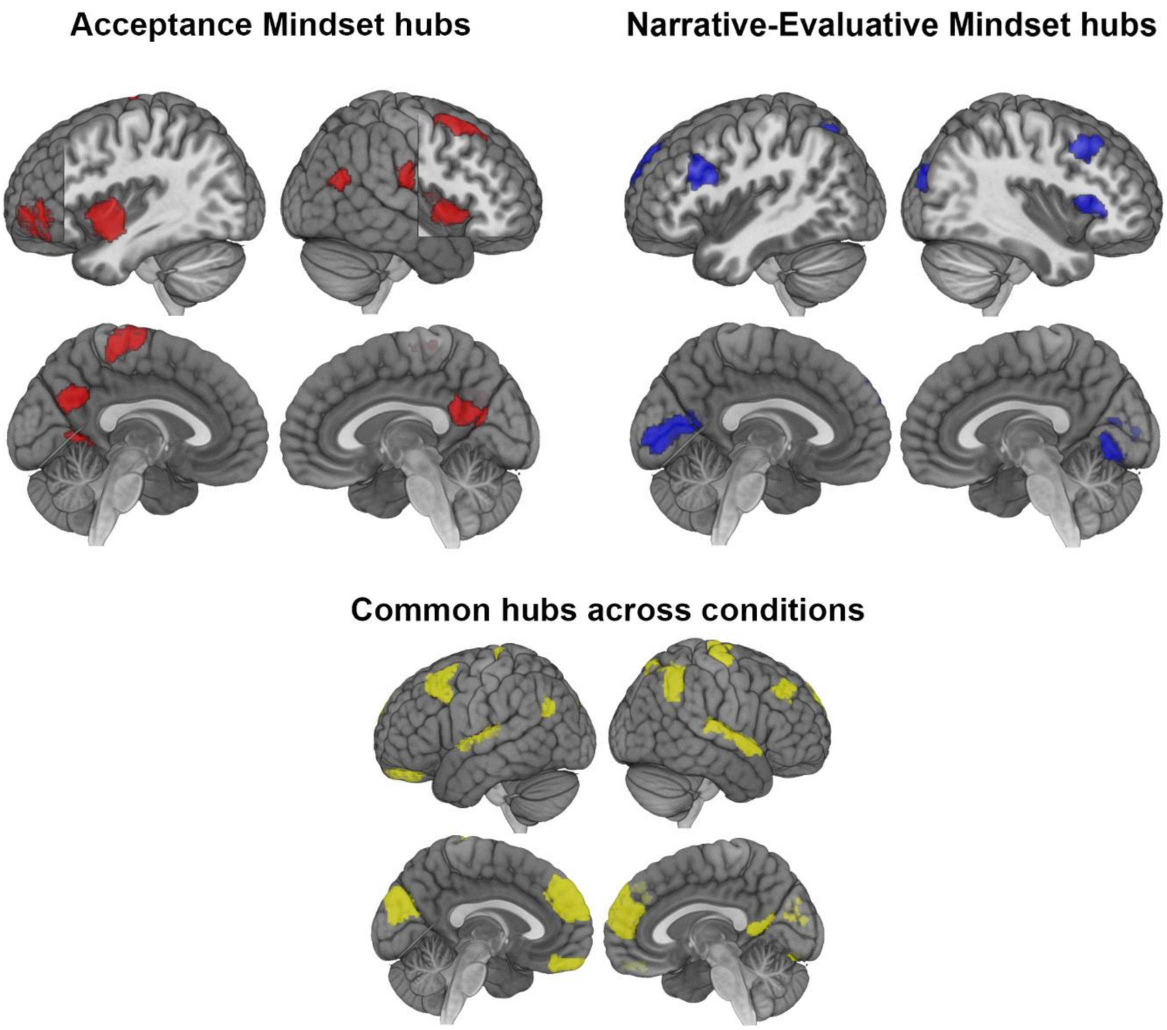


**Figure S3**. Provincial hubs during each condition. Provincial hubs have stronger within-network connections than other nodes in their home network, making them points of information flow and integration on a local level.

**Supplementary Methods**

### Questionnaires

Subjects also filled out a number of questionnaires immediately following the scanning session to place acceptance score in a broader context:

Five Facet Mindfulness Questionnaire (Baer, Smith, Hopkins, Krietemeyer, & Toney, 2006). This questionnaire measures the five facets that are thought to compose trait mindfulness: observing, describing, acting with awareness, non-judging of inner experience, and non-reactivity to inner experience. An example item is: "When I'm walking, I deliberately notice the sensations of my body moving". Each item is rated on a likert scale from 1 = never or very rarely true, to 5 = very often or always true. Scores are summed up for each individual category, as well as for overall.

Ruminative Response Scale-Brief (Treynor, Gonzalez, & Nolen-Hoeksema, 2003). This is a 22-item scale that is a shorter version of the original Ruminative Response Scale. Subjects indicate what they generally tend to think or do in response to depressive thoughts. An example item is: "Think about how sad you feel". Each item is rated on a likert scale from 1 = almost never, to 4 = almost always. Overall scores are summed up.

Big Five Inventory (John & Srivastava, 1999). This is a 44-item scale that measures an individual on the Big Five factors of personality (Extraversion, Agreeableness, Conscientiousness, Neuroticism, Openness). Example items include: "I see myself as someone who is talkative", or "I see myself as someone who is emotionally stable, not easily upset". Each item is rated on a likert scale from 1 = disagree strongly, to 5 = agree strongly. Scores are summed up for each individual category.

Trait Emotional Intelligence Questionnaire-Short Form (Petrides & Furnham, 2001). This is a 30-item scale is designed to measure global trait emotional intelligence. An example item is: "Expressing my emotions with words is not a problem for me". Each item is rated on a likert scale from 1 = strongly agree, to 7 = strongly disagree. Mean scores are calculated.

RiTE (ritualistic, theistic, and existential) measure of spirituality (Webb, Toussaint, & Dula, 2014). This scale consists of three 10-item subscales that each measure distinct aspects of spirituality: (1) Ritualistic Spirituality, (2) Theistic Spirituality, and (3) Existential Spirituality. Examples include: "I feel that helping others is very important", or "I believe in a deity or deities". Each item is rated on a likert scale from 1 = strongly disagree, to 5 = strongly agree. Overall scores are summed up.

Thought Control Questionnaire (Wells & Davies, 1994). This is a 30-item scale that assesses an individual's tendency to use strategies for the control of unpleasant and unwanted thoughts. The scale includes 5 subcategories, each of which represents a different strategy for thought control: distraction, social control, worry, punishment, and reappraisal. An example item is: "When I experience an unpleasant/unwanted thought, I call to mind positive images instead". Each item is rated on a likert scale from 1 = never, to 4 = almost always. Scores are summed up for each individual category, as well as for overall.

Santa Clara Brief Compassion Scale (Hwang, Plante, & Lackey, 2008). The Santa Clare Brief Compassion Scale is a brief 5-item index that assesses compassion and its link to prosocial behaviors. An example item is: "When I hear about someone (a stranger) going through a difficult time, I feel a great deal of compassion for him or her." Each item is rated on a likert scale from 1 = not at all true of me, to 7 = very true of me. Mean scores are calculated.

Satisfaction with Life Scale (Diener, Emmons, Larsen, & Griffin, 1985). This is a brief 5-item scale designed to measure global cognitive judgments of one's satisfaction with their life. An example item is: "In most ways, my life is close to my ideal". Each item is rated on a likert scale from 1 = strongly disagree, to 7 = strongly agree. Overall scores are summed up.

Flourishing Scale (Diener et al., 2010). This is a brief 8-item measure of the respondent's self-perceived success in important areas such as relationships, self-esteem, purpose, and optimism. The scale provides a single psychological well-being score. An example item is: "I lead a purposeful and meaningful life". Each item is rated on a likert scale from 1 = strongly disagree, to 7 = strongly agree. Overall scores are summed up.
